# TrustPGS: When can a polygenic score be trusted? A per-individual reliability framework across ancestries

**DOI:** 10.64898/2026.07.01.735762

**Authors:** Abdulmujeeb T. Onawole, Ridwanullahi A. Adegoke, Olalekan Amoo

## Abstract

Polygenic scores summarise genetic predisposition to a trait, but a population-level accuracy figure cannot tell a clinician whether a given prediction is reliable for the person in front of them. This gap is most consequential for individuals whose ancestry is under-represented in the discovery cohort, precisely the patients for whom a wrong trust call carries the highest clinical cost. We present TrustPGS, a framework that tells clinicians and downstream models which individual predictions can be trusted and which cannot, so that polygenic scores can inform clinical decisions rather than being acted on uniformly regardless of how well-supported each prediction is. The framework rests on two axes calibrated on a discovery cohort, the consensus of a Bayesian posterior-sample ensemble and the directional agreement of the top-magnitude linkage-disequilibrium blocks. We computed SBayesRC posterior-sample scores for ten polygenic traits in the 1000 Genomes Project phase-3 cohort and tested whether the resulting trust labels transfer, without recalibration, to the ancestrally diverse Simons Genome Diversity Project, comparing strict application of the European cutoffs, percentile-rank rescaling, and within-cohort recalibration. Percentile-rank rescaling preserved an enrichment factor above one in non-European populations for five of ten traits (Alzheimer disease, breast cancer, body mass index, LDL cholesterol, and systolic blood pressure), traits whose European and target-cohort distributions were shifted but comparable in shape. Three traits (coronary artery disease, height, and schizophrenia) carried distributions that differed in shape rather than location, a pattern traceable to discovery-cohort bias that recalibration could not repair either, and two further traits (type 2 diabetes and educational attainment) showed intermediate behaviour, present but never enriched in one case, and an apparent success that rank-mapping correctly unmasked as artefactual in the other. Because each of these patterns is detectable before any individual-level claim is made, TrustPGS gives clinicians and downstream models a falsifiable, per-trait basis for deciding when a reliability label can be trusted on a new population, rather than a single portability promise that holds or fails silently.

## 1. Introduction

Polygenic scores combine the small cumulative effects of many genetic variants into a single number estimating an individual’s predisposition to a trait. This predictive power is now accurate enough to be utilized in prospective clinical trials for risk-stratified screening, such as the WISDOM study for breast cancer (Eklund et al. 2018; Esserman 2017). The move from research into clinical use, however, exposes a gap that population-level summary statistics cannot close. Much of the ongoing work to maximize this predictive power focuses on population-level summary statistics, either by expanding the training cohorts horizontally across genetically correlated traits (Albiñana et al. 2023) or by integrating functional genomic annotations longitudinally across millions of variants (Zheng et al. 2024). Two individuals can receive identical PGS estimates while resting on very different evidence underneath, one supported by a large set of agreeing common variants, the other by a handful of rare and discordant ones (Ding et al. 2022). A clinician shown only the point estimate has no way to tell those two cases apart, and the stakes of that blind spot rise for any patient whose ancestry is poorly represented in the cohort the score was built on, since the population accuracy a PGS advertises was measured on people the patient may not resemble (Martin et al. 2019).

Existing work on PGS uncertainty has been dominated by Bayesian shrinkage methods that report a posterior on each variant’s effect (Ge et al. 2019; Privé et al. 2020), a posterior that can in principle be propagated forward into a posterior on the individual’s score. SBayesRC (Zheng et al. 2024) is the current best-in-class method in that family, returning a posterior mean and standard error for every variant across the genome. What is still missing, and what motivates this paper, is the next step, translating that variant-level posterior into a single label that a clinician or downstream model can use to decide which predictions to act on and which to treat with caution. Recent reviews of PGS interpretability identify this same gap, arguing that an explanation without an accompanying reliability tag is of limited use at the bedside (Schuran et al. 2025).

We address that gap with a two-axis labelling rule, adapting a general consensus-and-agreement trust framework whose foundations we established in evaluating explainable AI for molecular property prediction (Onawole et al. 2026) and validated as a four-scenario trust mechanism in an unrelated domain, attribution-channel agreement for predicting gas-separation performance in metal-organic frameworks (Onawole 2026). The second asks whether the genome regions contributing most to that score agree on its direction, which we call top-block attribution agreement. Neither axis is fully informative on its own under high polygenicity,, since polygenic prediction is by nature spread across many variants and a single discordant region does not make a prediction wrong. Combined, the two axes separate individuals whose prediction and underlying explanation both hold together from individuals where one or both diverge. We label the four resulting combinations Scenario A (consensus and agreement), Scenario B (consensus only), Scenario C (agreement only), and Scenario D (neither), calibrated at the 30th percentile of the consensus axis and the 70th percentile of the agreement axis on a discovery cohort, so the cutoffs are derived from data rather than asserted by convention. Figure 1 sets out the scheme.

**Figure 1.**
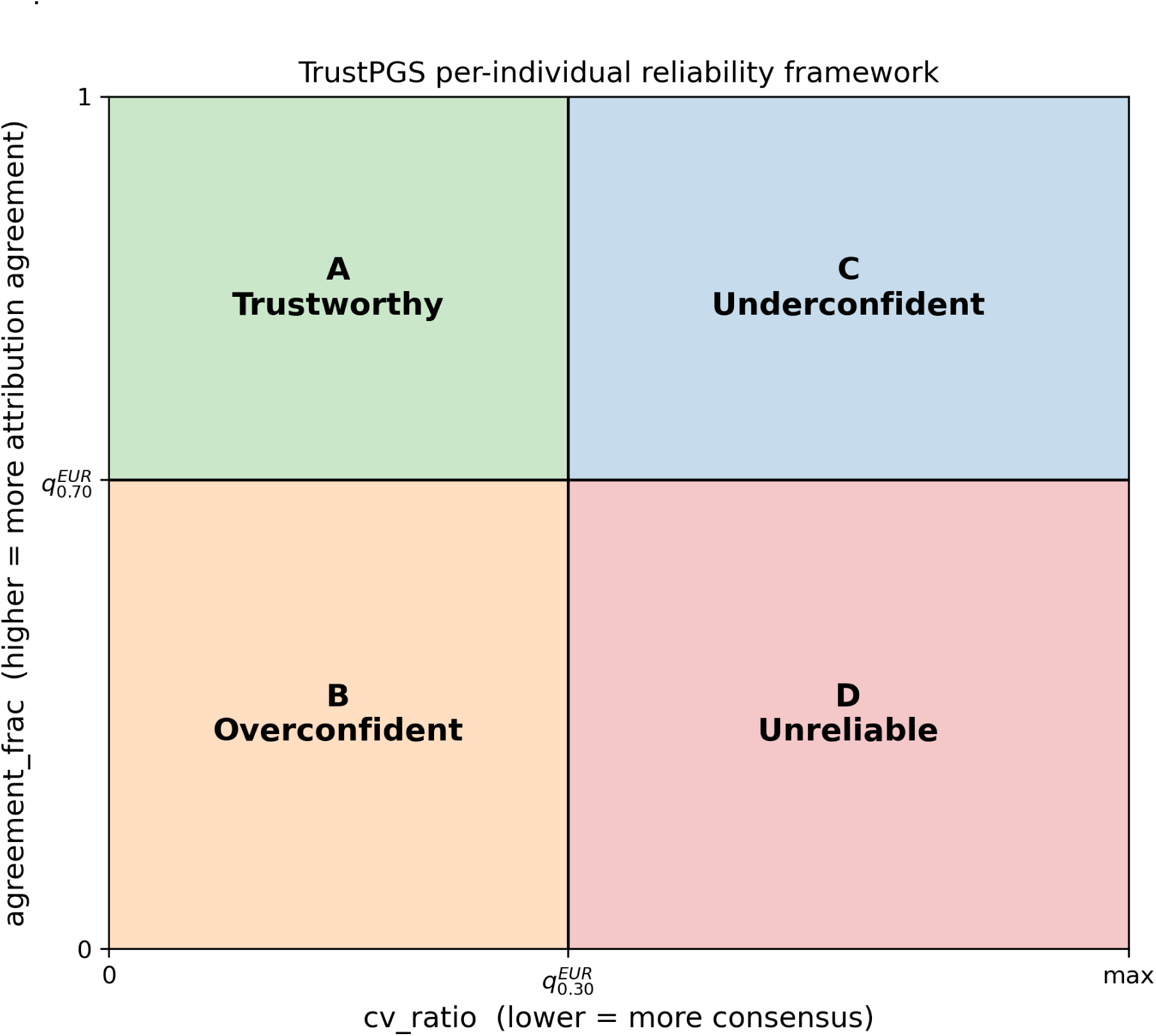
The TrustPGS per-individual trust framework. Each individual is positioned on two axes, ensemble consensus and top-block attribution agreement, with cutoffs calibrated on the 1000G EUR discovery fold dividing the plane into four quadrants. Scenario A (high consensus, high agreement) marks individuals whose prediction and explanation both transfer. Scenario D (low consensus, low agreement) marks the unreliable case. Scenarios B and C are the two asymmetric failure modes.

The question this raises for cross-ancestry use is direct, whether cutoffs calibrated on a European discovery cohort still identify a more reliable subset of individuals in non-European super populations (the broad continental-ancestry groupings, African, admixed American, East Asian, European, South Asian, that cohorts such as 1000 Genomes use to pool individually sampled populations) without being refitted. We set out three honest outcomes in advance of running the analysis. Full transfer would be a portability win for the framework as it stands. Partial transfer, with failures that can be diagnosed rather than merely observed, would show which trait architectures support a per-individual reliability claim outside the discovery population. Full failure would limit the contribution to within-cohort calibration only, with no cross-ancestry claim attached. We tested all three regimes, strict transfer, percentile-rank rescaling, and within-cohort recalibration, across the ten traits in our discovery panel. The result is partial transfer, and what distinguishes the traits that transfer from the traits that do not is itself diagnosable in advance, rather than only visible after the fact. The remainder of the paper sets out the framework and discovery-cohort behaviour in the Methods, tests transfer to the Simons Genome Diversity Project under the three calibration regimes, and shows how that diagnostic is built and what it reveals.

## 2. Methods

### 2.1 Cohorts and weight files

The discovery cohort is the 1000 Genomes Project phase-3 release (n=2504 across five super-populations), converted from VCF to PLINK2 binary format on a per-chromosome basis and filtered to the single nucleotide polymorphisms (SNPs) present in the SBayesRC weight files (Genomes Project 2015). The test cohort is the Simons Genome Diversity Project, using the cteam-extended release filtered to a minor allele frequency above 0.1 percent (n=345 across 142 populations) (Mallick et al. 2016). We map its populations onto the 1000G super-population scheme by continental region, with two exceptions, SouthAsia and CentralAsiaSiberia are both collapsed into SAS since neither has enough individuals to report separately, and Oceania is retained as a non-mapped category, OCN, since it has no 1000G analog. Sample sizes per super-population for both cohorts are given in Table 1. AMR (n=28) sits below the n>=30 enrichment reporting threshold, and OCN (n=26) is reported for completeness only.

**Table 1.** Sample size per 1000G super-population in each cohort. 1000G phase-3 is the discovery cohort and the source of the empirical-quantile cutoffs. SGDP is the strict-OOD test cohort. The * marks the two SGDP cells that are reported with caveat: AMR (n=28) is below the n>=30 enrichment-factor reporting threshold, and OCN (n=26) has no 1000G analog and is not used in the cross-ancestry transfer analysis.

| Cohort | n<br>total | AFR<br>(Africa) | AMR<br>(America) | EAS<br>(East<br>Asia) | EUR<br>(Europe) | SAS<br>(South<br>&<br>Central<br>Asia) | OCN<br>(Oceania) | Notes |
| --- | --- | --- | --- | --- | --- | --- | --- | --- |
| 1000G<br>phase-3 | 2504 | 661 | 347 | 504 | 503 | 489 | 0 | Discovery<br>cohort;<br>EUR fold<br>(n=503)<br>used to fit<br>empirical-quantile cutoffs. |
| SGDP | 345 | 93 | 28* | 48 | 75 | 75 | 26* | Strict-OOD test<br>cohort;<br>AMR<br>below<br>n>=30<br>reporting<br>threshold;<br>OCN has<br>no 1000G<br>analog. |

Per-SNP weights are taken from the SBayesRC 115-trait share, which provides a posterior mean effect on the A1 allele, a standard error, a posterior inclusion probability, and a per-iteration posterior sample stub for every SNP. All three data sources share one genome coordinate system, GRCh37 (also called hg19), so no liftover, the standard conversion between reference genome versions, was needed. We selected ten priority traits spanning three genetic architectures, highly polygenic quantitative traits (height, BMI (Body Mass Index), LDL(low-density lipoprotein) cholesterol), diseases with major-effect loci (Alzheimer disease, breast cancer), and psychiatric or behavioural traits (schizophrenia, educational attainment). The last category is named here in the field’s conventional statistical-genetics sense, any phenotype with a heritable component and a published GWAS, not as a claim of biological equivalence to the disease and quantitative traits in the panel. The full trait list and source GWAS are given in Table 2.

**Table 2.** Ten priority traits used in the present work. For each trait, the SBayesRC weight tarball, the variant-density set, the discovery population, and the source GWAS are given. All weight files are GRCh37/hg19; no liftover is performed because 1000G phase-3 and SGDP (hs37d5) are both GRCh37-compatible.

| Trait | SBayesRC file ID | Variant density | Discovery population | Source GWAS | Architecture note |
| --- | --- | --- | --- | --- | --- |

|  |  | set |  | (ref.) |  |
| --- | --- | --- | --- | --- | --- |
| Height | Height_03 | HD<br>(~7M) | EUR (UK Biobank +<br>GIANT) | (Yengo et<br>al. 2022) | Quantitative,<br>highly<br>polygenic |
| BMI<br>(Body Mass<br>Index) | BMI_03 | HD<br>(~7M) | EUR (UK Biobank +<br>GIANT) | (Yengo et<br>al. 2018) | Quantitative,<br>polygenic |
| T2D<br>(Type 2<br>Diabetes) | T2D_03 | HD<br>(~7M) | EUR (DIAMANTE) | (Mahajan et<br>al. 2022) | Disease,<br>polygenic |
| CAD<br>(Coronary<br>Artery<br>Disease ) | CAD_03 | HD<br>(~7M) | EUR<br>(CARDIoGRAMplusC4D) | (Aragam et<br>al. 2022) | Disease,<br>polygenic |
| Breast cancer | BreastCancer_03 | HD<br>(~7M) | EUR (BCAC) | (Michailidou<br>et al. 2017) | Disease,<br>includes<br>major BRCA<br>effects |
| Alzheimer<br>disease | Alzheimer_03 | HD<br>(~7M) | EUR (IGAP + UK<br>Biobank) | (Bellenguez<br>et al. 2022) | Disease,<br>dominated<br>by APOE |
| LDL<br>cholesterol | LDL_03 | HD<br>(~7M) | EUR (Global Lipids GC) | (Graham et<br>al. 2021) | Quantitative,<br>polygenic |
| Systolic blood<br>pressure<br>(SBP) | SBP_03 | HD<br>(~7M) | EUR (Evangelou ICBP) | (Evangelou<br>et al. 2018) | Quantitative,<br>polygenic |
| Schizophrenia | Schizophrenia_03 | HM3<br>(~1.16M) | EUR (PGC3 SCZ) | (Trubetskoy<br>et al. 2022) | Psychiatric,<br>polygenic |
| Educational<br>attainment | EA_03 | HM3<br>(~1.16M) | EUR (SSGAC EA4) | (Okbay et<br>al. 2022) | Behavioural,<br>highly<br>polygenic |

### 2.2 Posterior-sample ensemble PGS

For each trait and each individual, we draw K=20 posterior-sample effect vectors beta_k ∼ Normal(A1Effect, SE²) from SBayesRC’s per-SNP posterior moments, giving a posterior-sample PGS ensemble of K members per individual. The ensemble mean is the point PGS estimate. The ensemble standard deviation, std(pgs_ensemble), is the per-individual uncertainty in that score. We define cv_ratio = std(pgs_ensemble) / pop_std, where pop_std is the standard deviation of the ensemble-mean PGS across the discovery cohort. A low cv_ratio means the K draws agree on this individual’s score relative to how much PGS varies across the population, which we read as ensemble consensus. We use this posterior-sample ensemble rather than a bootstrap-over-SNPs ensemble we used in earlier development, because bootstrap variance reflects SNP-dropout randomness, not genuine Bayesian uncertainty, the two are not interchangeable for a consensus claim.

### 2.3 Top-block attribution and the agreement axis

Ensemble consensus, from 2.2, asks whether an individual’s score is stable across posterior draws. The agreement axis asks a different question, whether the genome regions driving that score agree with each other on its direction. For each individual and each ensemble member, the per-SNP contribution is the individual’s centred A1 dosage multiplied by the posterior-sample beta. We aggregate these into LD blocks using 1 Mb fixed bins, giving 2,697 blocks across chromosomes 1 to 22 for the SBayesRC high-density set, the simplest bin definition with no external dependency. For each individual and each block, the ensemble-mean contribution and its sign give the direction in which that block pushes the individual’s score away from the population mean. We take the top 10 percent of blocks by contribution magnitude per individual and compute agreement_frac, the fraction of those top blocks whose sign matches the individual’s overall deviation from the population mean. The 10 percent threshold was chosen after testing a range from 5 to 20 percent on the discovery cohort, 5 percent left too few blocks per individual for a stable statistic, and 20 percent diluted the signal by including blocks whose effects approximately cancel. Restricting to the top-magnitude blocks at all is essential, since asking every block to agree dilutes the signal under polygenicity and means the agreement axis rarely passes on most traits.

### 2.4 Empirical-quantile cutoffs and the four scenarios

Consensus and agreement are each a continuous number so far. Turning them into a decision means drawing a line on each and asking which side an individual falls on. Both cutoffs are calibrated on the 1000G EUR fold (n=503). The consensus cutoff is the 30th percentile of cv_ratio on EUR. The agreement cutoff is the 70th percentile of agreement_frac on EUR. Both are saved alongside the EUR pop_std and pop_mean, so downstream cohorts can inherit the rule without re-deriving it. An individual is Scenario A if cv_ratio is at or below the consensus cutoff and agreement_frac is at or above the agreement cutoff, Scenario B if only consensus passes, Scenario C if only agreement passes, and Scenario D otherwise. The quantiles are chosen so Scenario A is non-empty by construction on the calibration fold. The falsifiable claim is not whether 30 and 70 are the right percentiles, it is whether the labels they produce predict cross-ancestry reliability.

### 2.5 Three calibration regimes for the SGDP test

We compare three labelling regimes on SGDP. Regime 1, strict OOD, applies the 1000G EUR consensus cutoff, agreement cutoff, pop_std, and pop_mean unchanged. Regime 2, rank-mapped OOD, applies the cutoffs at the same percentile rank within SGDP’s own distributions (Table S1), the right test for the case where EUR and SGDP are shifted but similarly shaped. Regime 3, within-cohort recalibration, refits both cutoffs directly on the SGDP EUR fold (n=75), the weaker, cohort-own comparator. Each regime produces a Scenario label per individual and an enrichment factor per super-population.

### 2.6 Phenotype simulation and the enrichment factor

SGDP has no measured phenotype, so we simulate one. For each trait, y = g + noise, where g is the SBayesRC posterior-mean PGS and noise is drawn from Normal(0, var(g) × (1 − h²) / h²). We set h² = 0.4 for every trait, a mid-range heritability that avoids favouring any one trait’s true value, since SGDP individuals were never genotyped for any of these phenotypes and a trait-specific h² cannot be estimated from this data. The absolute prediction error, |pred − y|, is the per-individual reliability metric. The enrichment factor (EF) at the 0.5 SD threshold, EF@0.5, is the fraction of Scenario A individuals with error at or below 0.5 SD, divided by the same fraction across the whole super-population. EF@0.5 above one means Scenario A is more accurate than the population base rate; at or below one, the label carries no per-individual signal in that cell. We use 0.5 SD as the headline threshold because clinically actionable PGS decisions typically operate within a fraction of an SD of the population mean, and report 1.0 and 2.0 SD thresholds in the supplement. Full EF values at 0.5, 1.0, and 2.0 SD, for all three calibration regimes, all ten traits, and all four reportable super-populations, are given in Supporting Information Table S2.

## 3. Results

We report results in three stages. First, we confirm the framework behaves sensibly on the cohort it was calibrated on, before asking anything harder of it. Second, we test whether its labels transfer to a different ancestry without retraining, under three increasingly permissive calibration regimes, and find that they do for most traits, fail for a few in a way we can explain in advance, and behave in two further, distinct ways for the remaining two. Third, we discuss what that partial result means for deploying the framework, where it falls short, and how it relates to prior work.

### 3.1 Scenario A categorisation on the discovery cohort

On the 1000G discovery cohort itself, before any cross-ancestry claim is made, Scenario A needs to do two things to be worth using at all, it needs to actually contain people, and those people need to be more accurate than the population at large. Across the ten traits, Scenario A is populated for every super-population except AFR on five traits where the agreement axis is uniformly low (Figure 2), and EF@0.5 exceeds 1.0 in non-EUR super-populations on five traits, including height (EAS=1.29, SAS=1.27) and breast cancer (AFR=1.19, EAS=1.22, EUR=1.23). Three traits (CAD, Alzheimer, LDL) sit at or below EF=1.0 for most super-populations on 1000G, and these are the same traits that will fail or be rescued in different ways on the SGDP transfer test that follows, supporting the read that the framework is genuinely sensitive to trait architecture rather than labelling everyone the same. The sanity check r(pred, g) lies between 0.92 and 0.99 across every cell, confirming the underlying PGS computation is intact, so any failure of the trust labels that follows is a labelling failure, not a scoring one.

**Figure 2.**
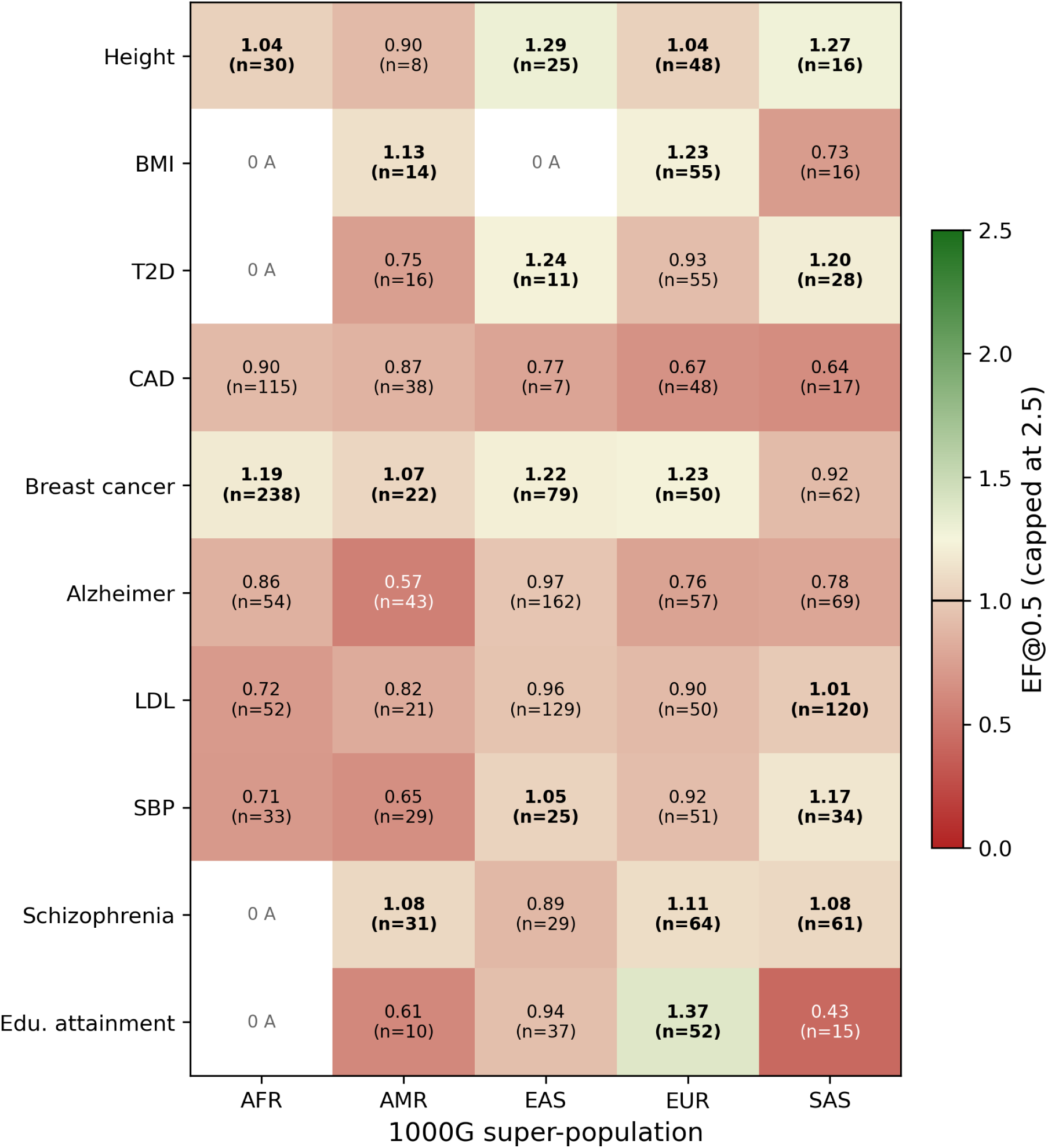
Enrichment factor at the 0.5 SD threshold (EF@0.5) for Scenario A on the 1000G discovery cohort, by trait and 1000G super-population. EF@0.5 > 1 means Scenario A precision exceeds the population base rate. Cells with empty Scenario A are marked 0 A. EUR is the calibration fold for the empirical-quantile cutoffs; non-EUR cells report transfer to other 1000G super-populations.

### 3.2 Trait performance transfer under strict cutoffs

Naively inheriting the 1000G EUR cutoffs onto SGDP gives a degenerate Scenario distribution on eight of ten traits (Table 3). Five traits collapse to all-D or all-B (BMI, CAD, LDL, SBP, T2D), because the 1000G EUR cv_ratio cutoff lands above the SGDP cv_ratio 95th percentile, so every SGDP individual passes the consensus axis, which combined with a strict agreement axis gives all-or-nothing labels. Two traits (height, alzheimer) show the opposite, the EUR cutoff sits below the SGDP 5th percentile, so almost nobody passes consensus. Schizophrenia shows a third failure mode, the EUR agreement_frac distribution lives in [0.86, 0.90] but the SGDP distribution lives in [0.09, 0.14], a clean and complete disjunction. Breast cancer (n_A=38) and educational attainment (n_A=55) are the only traits whose strict counts fall in a defensible middle range. The per-cohort sanity check r(pred, g) on SGDP stays in the 0.95 to 0.99 range for every trait, so the PGS itself transfers fine, but the trust label that does not.

**Table 3.** Number of SGDP individuals labelled Scenario A (consensus and agreement both pass) under each calibration regime, per trait. Differences across columns confirm that the percentile-rank and within-cohort calibration code paths fired; eight of ten traits change at least one regime. Two traits (CAD, schizophrenia) remain at A=0 under both strict and rank, matching the distributional diagnosis (Figure 4) that those two and height carry an EUR vs SGDP shape mismatch that location rescaling alone cannot repair.

| Trait | Strict OOD | Rank-mapped OOD | Recal (cohort-own) |
| --- | --- | --- | --- |
| Height | 0 | 0 | 50 |
| BMI | 12 | 25 | 16 |
| T2D | 4 | 19 | 8 |
| CAD | 0 | 0 | 36 |
| Breast cancer | 38 | 47 | 55 |
| Alzheimer disease | 4 | 178 | 25 |
| LDL cholesterol | 0 | 99 | 29 |
| Systolic blood | 288 | 153 | 14 |
| pressure |  |  |  |
| Schizophrenia | 0 | 0 | 11 |
| Educational attainment | 55 | 0 | 17 |

Figure 3 shows why the trust label does not transfer. Plotting the empirical cumulative distribution functions (CDFs, curves showing what fraction of a cohort falls at or below each value) of cv_ratio and agreement_frac for 1000G EUR and SGDP, per trait, with the inherited EUR cutoffs marked, separates the traits cleanly into two groups. For Alzheimer, BMI, and LDL, the EUR and SGDP CDFs sit side by side like translates of each other, a single linear shift in cv_ratio space would align them, which is exactly what the rank-mapping regime does next. For CAD, height, and schizophrenia, the CDFs differ in shape, not just location. Schizophrenia is the cleanest example, the SGDP agreement_frac CDF rises sharply in [0.05, 0.15] and is flat after that, while the EUR CDF rises in [0.85, 0.95], so no monotone rescaling brings the two cutoffs onto the same individuals. That the diagnostic separates rescuable from unrescuable traits before any rescaling code is even written is the strongest evidence the labels capture real per-trait structure, not a labelling artefact.

**Figure 3.**
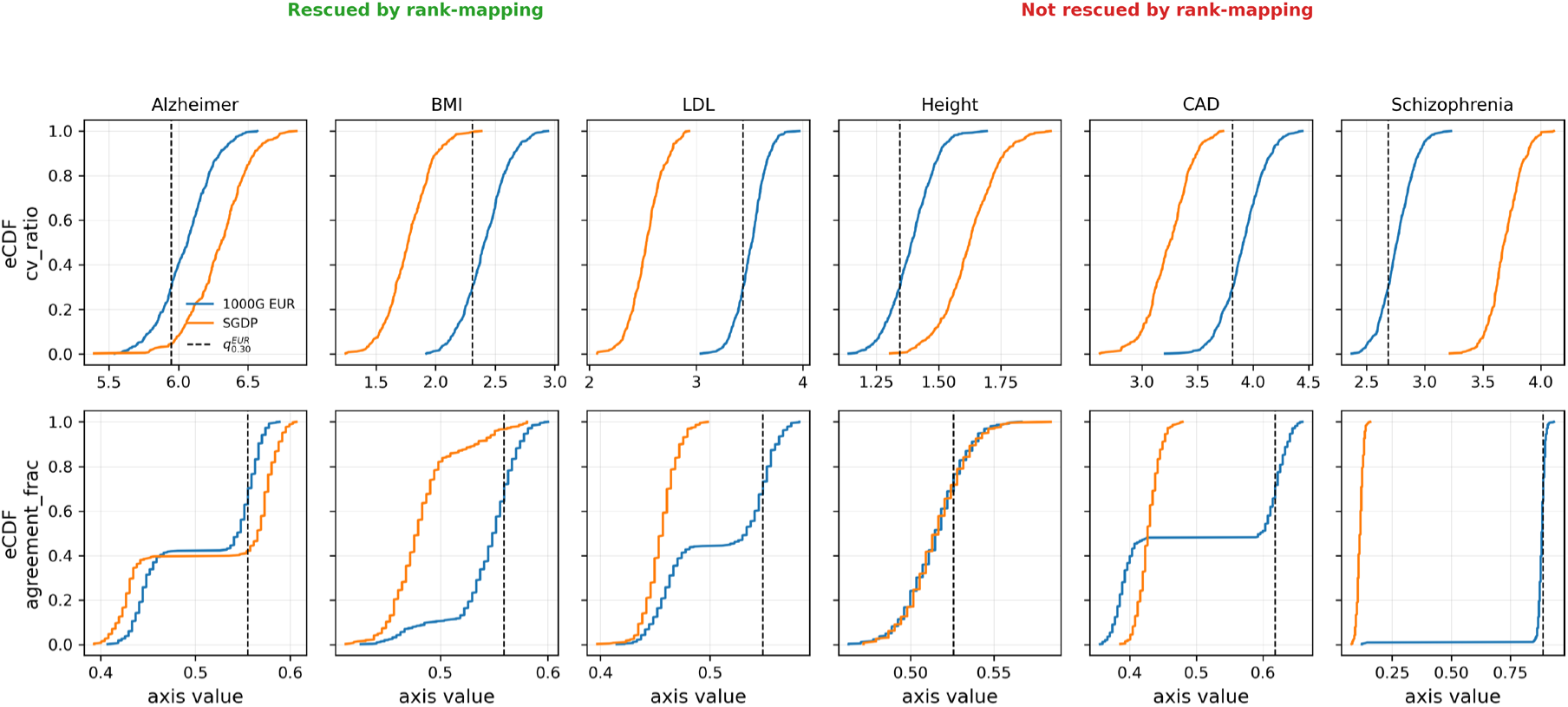
Empirical CDFs of the two trust axes for 1000G EUR (blue) and SGDP (orange), per trait. Top row, cv_ratio with the 1000G EUR 30th-percentile cutoff marked. Bottom row, agreement_frac with the 1000G EUR 70th-percentile cutoff marked. The three traits on the right (CAD, height, schizophrenia) are the ones not rescued by rank-mapping; their EUR and SGDP CDFs differ in shape rather than in location, which is the diagnostic signature of EUR-discovery bias that the framework exposes.

### 3.3 Trait performance transfer under rank-mapped cutoffs

Switching to the rank-mapped regime restores per-individual signal on the traits whose CDFs are shape-comparable but shifted. Table 4 gives the full grid, EF@0.5 exceeds 1.0 in EUR on five traits (Alzheimer 1.15, breast cancer 1.70, BMI 1.25, LDL 1.18, SBP 1.29), and in at least one non-European super-population on each of those five (Alzheimer SAS 1.24, breast cancer SAS 1.43, LDL EUR 1.18, SBP EUR 1.29, BMI SAS 1.01). Alzheimer is the most striking case, going from four Scenario A individuals under strict transfer to 178 under rank-mapping, with EF@0.5 above 1 in three of four super-populations, a result that only makes sense because the regime allows a location shift without touching the underlying rule. AFR is consistently the weakest super-population wherever it has enough individuals to report (n_A >= 5), falling below 1 on Alzheimer (0.47), breast cancer (0.72), and SBP (0.54), consistent with the broader PGS portability literature (Martin et al. 2019) and pointing to limits of the inherited European agreement axis when applied to predominantly African individuals.

**Table 4.** Enrichment factor at the 0.5 SD threshold (EF@0.5) under the rank-mapped OOD regime (regime 2), per trait and per SGDP super-population, with the number of Scenario A individuals contributing to each estimate (n_A). Cells with n_A < 5 are shown in parentheses with the symbol † to flag that the estimate is statistically unstable. 0 A indicates an empty Scenario A cell. EUR is shown for completeness; the cross-ancestry transfer comparison is in the non-EUR columns.

| Trait | AFR | EAS | EUR | SAS |
| --- | --- | --- | --- | --- |
| Height | 0 A | 0 A | 0 A | 0 A |
| BMI | (1.33)† $n_A=2$ | 0 A | 1.25, $n_A=12$ | 1.01, $n_A=9$ |
| T2D | 0 A | (0.00)† $n_A=1$ | 0.88, $n_A=15$ | (0.00)† $n_A=3$ |
| CAD | 0 A | 0 A | 0 A | 0 A |
| Breast cancer | 0.72, $n_A=17$ | 1.37, $n_A=5$ | 1.70, $n_A=11$ | 1.43, $n_A=10$ |
| Alzheimer disease | 0.47, $n_A=14$ | 1.03, $n_A=34$ | 1.15, $n_A=56$ | 1.24, $n_A=51$ |
| LDL cholesterol | 1.16, n_A=5 | 0.86, n_A=23 | 1.18, n_A=22 | 0.96, n_A=28 |
| Systolic blood pressure | 0.54, n_A=19 | 0.91, n_A=30 | 1.29, n_A=38 | 1.17, n_A=38 |
| Schizophrenia | 0 A | 0 A | 0 A | 0 A |
| Educational attainment | 0 A | 0 A | 0 A | 0 A |

CAD, height, and schizophrenia remain at zero Scenario A individuals under both strict and rank-mapped regimes, exactly the three traits whose CDFs differed in shape in Figure 3, confirming that percentile-rank mapping cannot rescue a shape mismatch, only a shift. Figure 4 (middle panel) shows this directly, the same three traits stay empty across every regime tested. Schizophrenia shows the mechanism most clearly, the top-magnitude LD blocks driving the EUR score carry directions that do not transfer to SGDP individuals, so the EUR-derived agreement cutoff lands above almost every SGDP value. This is the per-individual analog of the well-documented EUR-discovery bias for psychiatric traits, but expressed as a flag on individual unreliability rather than a population-level accuracy drop. We read CAD and height as the same mechanism in non-psychiatric traits, where the top-block directions reflect EUR-cohort artefacts that the diagnostic catches automatically. T2D and educational attainment fit neither group and are treated separately.

**Figure 4.**
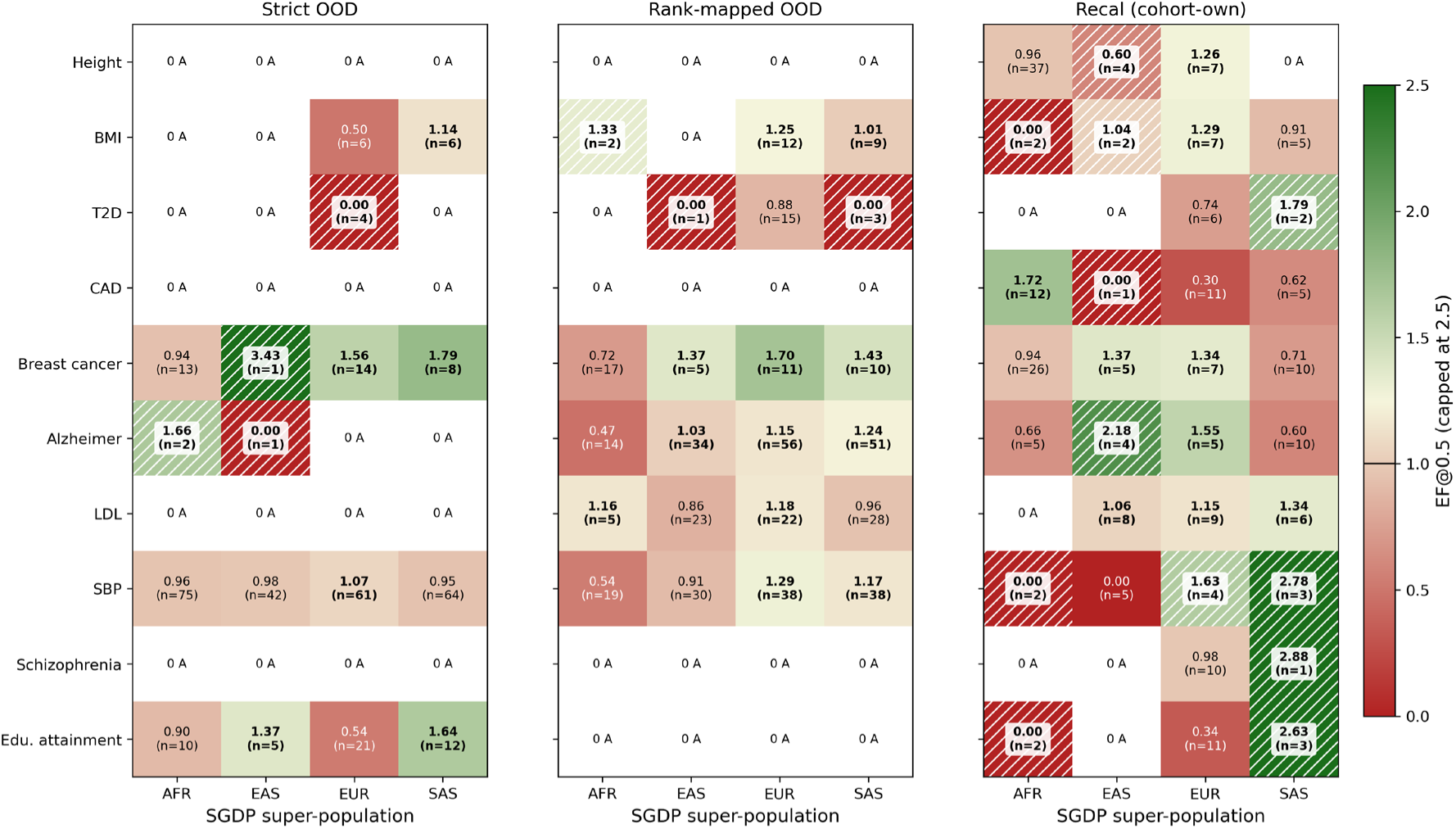
Cross-ancestry transfer of TrustPGS labels to the SGDP cohort under three calibration regimes. Strict OOD inherits 1000G EUR cutoffs and the EUR pop_std and pop_mean unchanged. Rank-mapped OOD inherits the EUR empirical distributions of cv_ratio and agreement_frac and applies the cutoffs at the same percentile rank on SGDP, while using cohort-own pop_std. Within-cohort recalibration refits both cutoffs on the SGDP EUR fold (n=75). Cells with n_A < 5 are hatched to flag that the enrichment estimate is statistically unstable.

### 3.4 Analysing the intermediate cases: T2D and educational attainment

T2D and educational attainment sit between the five rescued traits and the three shape-mismatch traits, each in a different way. T2D produces Scenario A individuals under every calibration regime (strict transfer 4, rank-mapped transfer 19, within-cohort recalibration 8, Table 3), but EF@0.5 under rank-mapping is 0.88 in EUR (Table 4), the only super-population with enough individuals to assess, so T2D produces a trust label that is not backed by any real accuracy gain.

Educational attainment shows the opposite, and more informative, pattern. Under strict transfer it produces 55 Scenario A individuals, a count comparable to breast cancer’s 38, but under rank-mapping that count collapses entirely to zero (Table 3). The collapse runs in the opposite direction from every rescued trait and from T2D, and the mechanism is artefactual strict-transfer labelling, the 1000G EUR cv_ratio cutoff happened to intersect the raw SGDP cv_ratio distribution in a narrow window by coincidence, and rank-mapping removes that accidental alignment by comparing through the EUR empirical percentile distribution rather than raw values. The strict-versus-rank discrepancy here is therefore a demonstration of the diagnostic working correctly, not a failure of it, the framework catches its own false positive. We treat T2D as a sub-threshold signal case and educational attainment as a masked artefact case.

### 3.5 Trait performance transfer under within-cohort recalibration

Recalibrating directly on SGDP does not rescue the three failing traits, it redistributes their labels without making them more accurate. Refitting both cutoffs directly on the SGDP EUR fold produces non-empty Scenario A counts for every trait (Table 3, recalibration column), but the EF pattern on the traits that already failed under strict and rank transfer is erratic. On CAD, recalibration gives 36 Scenario A individuals, but EUR EF=0.30 and AFR EF=1.72, the trait sits below population base rate in the exact super-population the cutoffs were fit on. On height, recalibration gives 50 Scenario A individuals with EUR EF=1.26, AFR EF=0.96 (no enrichment), and SAS EF=0.00, a pattern that doesn’t read as evidence of useful labels. On schizophrenia, recalibration gives 11 Scenario A individuals with EUR EF=0.98 and one SAS individual driving a spurious EF=2.88. Sample size is the limiting factor throughout, SGDP EUR is n=75, too small to fit two empirical-quantile cutoffs cleanly, so recalibration mostly reshuffles labels rather than recovering signal where strict and rank transfer both already failed.

## 4. Discussion

Per-individual reliability labels calibrated on a single ancestry do not transfer uniformly to a new one, but where they fail is predictable rather than arbitrary. Five of ten traits transfer cleanly under rank-mapped rescaling, their between-cohort distributions are shape-comparable, just shifted. Three traits fail outright (CAD, height, schizophrenia), their distributions differ qualitatively in shape, not merely in location, and no rescaling regime recovers them. The remaining two traits, T2D and educational attainment, behave in two further, distinct intermediate ways. This three-way split matters because it shows the framework is not simply unreliable outside its discovery population, it is informatively unreliable, the same diagnostic that flags which traits transfer also flags which traits will not, before any individual-level claim is made on them. That distinction, between failing silently and failing legibly, is what TrustPGS is meant to deliver.

A label that fails legibly is more useful than one that fails silently. It lets a downstream user run the rank-mapped regime on a new cohort, inspect the CDF overlay, and decide, before making any individual-level claim, whether a trait sits in the rescuable or the shape-mismatched regime. That is a stronger position than a single headline EF number, since it hands the user a falsifiable, per-trait diagnostic rather than a portability promise that holds or fails silently. We caution against reading the three failing traits as a condemnation of the underlying PGS itself, the sanity check r(pred, g) stays high for all three, so the failure belongs to the trust label, not the score, and is best attributed to EUR-discovery bias already documented in the underlying SBayesRC weights for those traits (Martin et al. 2019).

This interpretation carries three limitations worth weighing against it. First, the empirical-quantile cutoffs are fit on the whole 1000G EUR fold rather than a held-out split, so EUR-on-EUR EF estimates carry a small in-sample leak. We report the in-cohort numbers (Figure 2) for completeness, but the cross-ancestry claim does not depend on the EUR column. Second, the ensemble draws K=20 posterior samples from per-SNP marginal standard errors rather than the joint posterior, which would require running SBayesRC’s MCMC end-to-end, so the cv_ratio axis under-represents between-SNP correlation in the uncertainty. If anything, this makes Scenario A more permissive and biases EF estimates downward, so it is a conservative limitation rather than one that inflates our results. Third, SGDP’s sample size per super-population is small (n=47 to 93 in the four reportable cells), limiting the precision of every EF estimate, cells with n_A < 5 are marked unstable in Table 4 and Figure 3, but a larger SGDP-equivalent cohort would sharpen the interpretation throughout.

Set against prior work, this contribution is also clearly bounded. Prior work on PGS interpretability has focused on the model side, post-hoc attribution of deep-learning PRS through integrated gradients or SHAP, or feature-importance analysis of penalised-regression PRS (Lundberg and Lee 2017; Sundararajan et al. 2017). These approaches have advanced what an explanation can show, but none establish whether the explanation itself is reliable enough to act on, so they do not address which individual predictions a clinician or downstream model should trust. Prior work on PGS has increasingly focused on individual-level uncertainty, establishing rigorous frameworks to bound the inherent estimation and prediction errors of polygenic scores for a given individua (Ding et al. 2022; Sun et al. 2021; Wang et al. 2025). That body of work is the direct input to our framework, but it stops at a continuous interval rather than a per-individual decision rule. The contribution of TrustPGS is to combine an ensemble-consensus axis with a directional-agreement axis on the same individual, and to expose the cross-ancestry transfer of the resulting rule as a per-trait diagnostic, which neither line of prior work tests.

## 5. Conclusion

TrustPGS labels every PGS prediction with a per-individual reliability tag derived from two empirically calibrated axes, the ensemble consensus of a posterior-sample PGS and the directional agreement of the top-magnitude LD blocks. On 1000G phase-3 the framework is non-degenerate and shows trait-dependent behaviour rather than uniform labels. On SGDP it transfers across ancestries without recalibration on five of ten priority traits under percentile-rank rescaling, shows intermediate behaviour on two further traits (T2D and educational attainment), and on the three traits where it does not transfer at all (CAD, height, schizophrenia) the failure is diagnosable as a distribution-shape mismatch that within-cohort recalibration also fails to repair. The framework therefore gives a clinician or downstream model both a per-individual trust tag where applicable and a per-trait warning where the tag should not be trusted, a more honest deployment posture than a single population-level performance number. Two limitations bound how far this can be generalised before validation on a larger, independent non-European cohort, the small SGDP sample size per super-population, and an ensemble built from a marginal rather than joint posterior. Future work should extend the ensemble to multiple methods (SBayesRC, LDpred2, PRS-CS) so that the consensus axis captures cross-method as well as within-method uncertainty, and should validate the diagnostic on additional non-European cohorts (All of Us, H3Africa, GenomeAsia 100K) where the per-individual reliability question is most clinically pressing.

## Data and code availability

Code available here: https://github.com/MujeebOnawole/TrustPGS

## Acknowledgements

This work was supported by resources provided by The University of Queensland Research Computing Centre’s Bunya supercomputer.

