## Supplementary material for "TrustPGS: When can a polygenic score be trusted? A per-individual reliability framework across ancestries": Supplemntal Information

### S1. Extended methods

#### *S1.1 Compute environment and reproducibility*

All analyses were run inside a fixed Apptainer container providing Python 3.9, PyTorch, NumPy, SciPy, pandas, and scikit-learn. PLINK2 v2.0 alpha-6 (2025-01-29) was deployed as a static binary alongside the pipeline; additional Python packages were installed with pip --no-deps to avoid overriding the container's compiled dependencies. The container was treated as read-only throughout. The full pipeline is reproducible from the public 1000G phase-3 release, the public SGDP cteam_extended.v4 release, and the public SBayesRC 115-trait share, plus the source code in the TrustPGS repository.

#### *S1.2 Full SBayesRC weight file specification*

Each SBayesRC trait file is a whitespace-separated table with columns SNP (rsID), Chr, Pos, A1, A2, A1Freq, BETA (posterior mean effect on A1), SE, PIP (posterior inclusion probability), and BETAlast (the per-iteration MCMC sample stub used for posterior-draw reconstruction). The high-density variant set contains 7,356,518 SNPs; the HapMap3 fallback set contains 1,159,648 SNPs; we use the HD set for height, BMI, T2D, CAD, breast cancer, Alzheimer disease, LDL, and SBP and the HM3 set for schizophrenia and educational attainment, following the SBayesRC share defaults. All build coordinates are GRCh37/hg19, matching 1000G phase-3 and SGDP (hs37d5), so no liftover is performed.

#### *S1.3 LD-block definition and TOP_BLOCKS_FRACTION sensitivity*

LD blocks are defined as 1 Mb fixed bins over chr1-22, giving 2,697 blocks in the HD set. We chose fixed bins over the LDetect approach because fixed bins have no external dependency and the results are robust to bin size in the 0.5 to 2 Mb range (Spearman rho >= 0.95 between Scenario A label assignments at 1 Mb and 2 Mb on 1000G EUR for height). TOP_BLOCKS_FRACTION was set to 0.10 after a small sweep on the 1000G EUR fold for height; fractions in [0.05, 0.20] all produced non-degenerate Scenario A populations, with 0.05 producing too few blocks per individual to support a stable agreement statistic (sensitivity to one or two outlier blocks) and 0.20 diluting the signal by including blocks whose effects approximately cancel.

#### *S1.4 Sanity check on Bayesian posterior-sample ensemble*

Each ensemble member is drawn as beta_k ~ Normal(A1Effect, SE^2) using the SBayesRC per-SNP posterior mean and SE. We use K=20 members per trait and per individual; cv_ratio estimates stabilise at K>=15 in our 1000G EUR sensitivity tests on height (variance of cv_ratio across K from 15 to 40 is <2 percent of cv_ratio itself), so K=20 is a defensible middle ground between cost and stability.

#### *S1.5 Computational pipeline overview*

The pipeline is implemented as eight steps (download, extract, compute PGS, build tensor, simulate phenotype, ensemble PGS, explanation and labelling, final evaluation), chained by scheduler dependencies. Each step reads exclusively from the previous step's output and writes to a step-specific directory, so a failed step can be restarted in isolation without rerunning the full pipeline.

### S2. Two methodological corrections to cross-cohort label inheritance

Two implementation issues identified during cross-cohort validation are documented here because their uncorrected state would have produced misleading strict-OOD numbers; the results reported in the main manuscript are post-correction. **Table S1** summarises the effects and remedies; the description below provides context for readers seeking to replicate the diagnostic.

**Table S1.** Provenance of two cross-cohort corrections that materially shaped the strict-OOD readout, documented here so the reported CAD, height, and schizophrenia failure can be distinguished from a residual implementation artefact. Fix 19 corrected a population-mapping bug that left the SGDP per-super-population block empty for all ten traits. Fix 20 corrected a cohort-dependence bug in the trust-cutoff calculation that placed the inherited 1000G EUR cutoff entirely outside the SGDP cv_ratio range for five traits, producing uninterpretable all-A or all-B label distributions. All results reported in the main manuscript are post-fix.

| Fix # | Symptom (pre-fix) | Effect on result (pre-fix) | Fix | Result (post-fix) |
|---|---|---|---|---|
| 19 | utils.read_sgdp_panel keyed on metadata | Every SGDP individual | Route IID -> .fam FID (population) -> metadata | Strict-OOD: |

| | | | | |
|---|---|---|---|---|
| | sample-ID columns; xai.npz samples carry .fam IID instead | mapped to UNK -> by_super_pop block empty for all traits | Region -> SGDP_REGION_TO_SUPER | SGDP per-super-pop EF reported (4 super-pops covered) |
| 20 | cv_ratio = pgs_std / pop_std and direction = sign(pgs_mean - pgs_mean.mean()) both depend on cohort-own pop_std and pgs_mean | 1000G EUR cutoff landed entirely outside the SGDP cv_ratio range for 7 of 10 traits, giving bimodal all-A or all-B Scenario distributions | Freeze 1000G EUR pop_std and pop_mean into xai.npz; xai.run_trait raises if those scalars are absent on the OOD run | Strict-OOD: cv_ratio is now cohort-invariant; failure mode shifts from scale mismatch to shape mismatch (the diagnostic motivating regimes 2 and 3) |

The first correction addressed a population-mapping issue. The SGDP individual identifiers in the genotype processing output follow the cteam_extended FAM file convention, which differs from the identifier scheme used in the SGDP metadata sample columns (Sample_ID, Illumina_ID, SGDP_ID). The original mapping function keyed the lookup on the metadata columns and therefore failed to match any individual, so every SGDP sample mapped to an unknown population and the per-super-population evaluation output was empty for all ten traits. The correction routes each individual identifier through the FAM file population field to the metadata Region column and then to the 1000G super-population scheme, and raises an explicit error if the FAM path is absent rather than silently returning an empty result.

The second correction addressed a cohort-dependence issue in the trust-labelling step. The original definition cv_ratio = pgs_std / pop_std uses cohort-own pop_std, so the cv_ratio value for the same individual changes if the same SBayesRC weights are re-applied on a different cohort with a different population spread of pgs_mean. Because SGDP includes 142 populations versus 1000G's five EUR populations, the SGDP-derived pop_std differs substantially from the 1000G EUR pop_std for several traits, which placed the inherited 1000G EUR cv_ratio cutoff entirely outside the SGDP cv_ratio range for five traits (BMI, CAD, LDL, SBP, T2D) and produced bimodal all-A or all-B Scenario distributions that were not interpretable. The fix freezes pop_std and pop_mean from the source cohort run into each xai .npz, makes trust_labels() accept pop_std_override and pop_mean_override, and routes the inheritance through the --load-cutoffs-from flag so the SGDP run uses the EUR values unchanged. Without this fix the strict-OOD failure mode would have been a scale mismatch artefact; with the fix the failure mode is a genuine distribution-shape mismatch (Figure 4).

***S2.1 Phenotype simulation and the choice of h2***

SGDP carries no phenotype, so for the cross-ancestry evaluation we simulate y_i = g_i + eps_i per individual and per trait, where g_i is the SBayesRC posterior-mean PGS for individual i and eps_i is drawn from Normal(0, var(g)*(1-h2)/h2) with h2 = 0.4 across traits. h2 = 0.4 is a deliberately conservative middle estimate that does not favour any particular trait architecture; the absolute prediction error |pred - y| scales linearly with h2 in the sense that EF@0.5 SD is invariant under any monotone rescaling of h2 because both numerator and denominator share the same scale. The simulated phenotype was sanity-checked by computing Pearson r(pred, g) on each cohort and confirming the 0.92 to 0.99 range across every trait and super-population.

**Table S2.** Full per-trait, per-super-population enrichment factor at the 0.5, 1.0, and 2.0 SD thresholds, under all three calibration regimes. n_scenario_a is the count of Scenario A individuals contributing to the enrichment estimate at each threshold; base_rate is the fraction of all SGDP individuals in the super-population whose absolute prediction error is within the threshold; scenario_a_precision is the same fraction restricted to Scenario A; EF is the ratio scenario_a_precision / base_rate. The main-text Table 4 is the threshold=0.5, regime=rank slice.

| regime | trait | super_pop | n_pop | threshold_SD | n_scenario_a | base_rate | scenario_a_precision | EF |
|---|---|---|---|---|---|---|---|---|
| strict | Height | AFR | 93 | 0.5 | 0 | 0.452 | 0.000 | 0.000 |
| strict | Height | AFR | 93 | 1.0 | 0 | 0.742 | 0.000 | 0.000 |
| strict | Height | AFR | 93 | 2.0 | 0 | 0.957 | 0.000 | 0.000 |
| strict | Height | EAS | 48 | 0.5 | 0 | 0.417 | 0.000 | 0.000 |

| | | | | | | | | |
|---|---|---|---|---|---|---|---|---|
| strict | Height | EAS | 48 | 1.0 | 0 | 0.708 | 0.000 | 0.000 |
| strict | Height | EAS | 48 | 2.0 | 0 | 0.958 | 0.000 | 0.000 |
| strict | Height | EUR | 75 | 0.5 | 0 | 0.453 | 0.000 | 0.000 |
| strict | Height | EUR | 75 | 1.0 | 0 | 0.747 | 0.000 | 0.000 |
| strict | Height | EUR | 75 | 2.0 | 0 | 0.973 | 0.000 | 0.000 |
| strict | Height | SAS | 75 | 0.5 | 0 | 0.387 | 0.000 | 0.000 |
| strict | Height | SAS | 75 | 1.0 | 0 | 0.653 | 0.000 | 0.000 |
| strict | Height | SAS | 75 | 2.0 | 0 | 0.960 | 0.000 | 0.000 |
| strict | BMI | AFR | 93 | 0.5 | 0 | 0.376 | 0.000 | 0.000 |
| strict | BMI | AFR | 93 | 1.0 | 0 | 0.591 | 0.000 | 0.000 |
| strict | BMI | AFR | 93 | 2.0 | 0 | 0.925 | 0.000 | 0.000 |
| strict | BMI | EAS | 48 | 0.5 | 0 | 0.479 | 0.000 | 0.000 |
| strict | BMI | EAS | 48 | 1.0 | 0 | 0.667 | 0.000 | 0.000 |
| strict | BMI | EAS | 48 | 2.0 | 0 | 0.875 | 0.000 | 0.000 |
| strict | BMI | EUR | 75 | 0.5 | 6 | 0.333 | 0.167 | 0.500 |
| strict | BMI | EUR | 75 | 1.0 | 6 | 0.600 | 0.333 | 0.556 |

| | | | | | | | | |
|---|---|---|---|---|---|---|---|---|
| strict | BMI | EUR | 75 | 2.0 | 6 | 0.933 | 1.000 | 1.071 |
| strict | BMI | SAS | 75 | 0.5 | 6 | 0.440 | 0.500 | 1.136 |
| strict | BMI | SAS | 75 | 1.0 | 6 | 0.733 | 0.500 | 0.682 |
| strict | BMI | SAS | 75 | 2.0 | 6 | 0.947 | 0.667 | 0.704 |
| strict | T2D | AFR | 93 | 0.5 | 0 | 0.258 | 0.000 | 0.000 |
| strict | T2D | AFR | 93 | 1.0 | 0 | 0.559 | 0.000 | 0.000 |
| strict | T2D | AFR | 93 | 2.0 | 0 | 0.892 | 0.000 | 0.000 |
| strict | T2D | EAS | 48 | 0.5 | 0 | 0.417 | 0.000 | 0.000 |
| strict | T2D | EAS | 48 | 1.0 | 0 | 0.667 | 0.000 | 0.000 |
| strict | T2D | EAS | 48 | 2.0 | 0 | 0.833 | 0.000 | 0.000 |
| strict | T2D | EUR | 75 | 0.5 | 4 | 0.227 | 0.000 | 0.000 |
| strict | T2D | EUR | 75 | 1.0 | 4 | 0.493 | 0.000 | 0.000 |
| strict | T2D | EUR | 75 | 2.0 | 4 | 0.800 | 0.250 | 0.312 |
| strict | T2D | SAS | 75 | 0.5 | 0 | 0.280 | 0.000 | 0.000 |
| strict | T2D | SAS | 75 | 1.0 | 0 | 0.573 | 0.000 | 0.000 |
| strict | T2D | SAS | 75 | 2.0 | 0 | 0.880 | 0.000 | 0.000 |

| | | | | | | | | |
|---|---|---|---|---|---|---|---|---|
| strict | CAD | AFR | 93 | 0.5 | 0 | 0.290 | 0.000 | 0.000 |
| strict | CAD | AFR | 93 | 1.0 | 0 | 0.591 | 0.000 | 0.000 |
| strict | CAD | AFR | 93 | 2.0 | 0 | 0.925 | 0.000 | 0.000 |
| strict | CAD | EAS | 48 | 0.5 | 0 | 0.229 | 0.000 | 0.000 |
| strict | CAD | EAS | 48 | 1.0 | 0 | 0.625 | 0.000 | 0.000 |
| strict | CAD | EAS | 48 | 2.0 | 0 | 0.875 | 0.000 | 0.000 |
| strict | CAD | EUR | 75 | 0.5 | 0 | 0.307 | 0.000 | 0.000 |
| strict | CAD | EUR | 75 | 1.0 | 0 | 0.533 | 0.000 | 0.000 |
| strict | CAD | EUR | 75 | 2.0 | 0 | 0.867 | 0.000 | 0.000 |
| strict | CAD | SAS | 75 | 0.5 | 0 | 0.320 | 0.000 | 0.000 |
| strict | CAD | SAS | 75 | 1.0 | 0 | 0.547 | 0.000 | 0.000 |
| strict | CAD | SAS | 75 | 2.0 | 0 | 0.867 | 0.000 | 0.000 |
| strict | Breast cancer | AFR | 93 | 0.5 | 13 | 0.409 | 0.385 | 0.941 |
| strict | Breast cancer | AFR | 93 | 1.0 | 13 | 0.624 | 0.538 | 0.863 |
| strict | Breast cancer | AFR | 93 | 2.0 | 13 | 0.871 | 0.769 | 0.883 |
| strict | Breast cancer | EAS | 48 | 0.5 | 1 | 0.292 | 1.000 | 3.429 |

| | | | | | | | | |
|---|---|---|---|---|---|---|---|---|
| strict | Breast cancer | EAS | 48 | 1.0 | 1 | 0.583 | 1.000 | 1.714 |
| strict | Breast cancer | EAS | 48 | 2.0 | 1 | 0.917 | 1.000 | 1.091 |
| strict | Breast cancer | EUR | 75 | 0.5 | 14 | 0.320 | 0.500 | 1.562 |
| strict | Breast cancer | EUR | 75 | 1.0 | 14 | 0.627 | 0.571 | 0.912 |
| strict | Breast cancer | EUR | 75 | 2.0 | 14 | 0.933 | 0.929 | 0.995 |
| strict | Breast cancer | SAS | 75 | 0.5 | 8 | 0.280 | 0.500 | 1.786 |
| strict | Breast cancer | SAS | 75 | 1.0 | 8 | 0.547 | 0.500 | 0.915 |
| strict | Breast cancer | SAS | 75 | 2.0 | 8 | 0.880 | 1.000 | 1.136 |
| strict | Alzheimer disease | AFR | 93 | 0.5 | 2 | 0.301 | 0.500 | 1.661 |
| strict | Alzheimer disease | AFR | 93 | 1.0 | 2 | 0.591 | 0.500 | 0.845 |
| strict | Alzheimer disease | AFR | 93 | 2.0 | 2 | 0.903 | 1.000 | 1.107 |
| strict | Alzheimer disease | EAS | 48 | 0.5 | 1 | 0.229 | 0.000 | 0.000 |
| strict | Alzheimer disease | EAS | 48 | 1.0 | 1 | 0.479 | 0.000 | 0.000 |
| strict | Alzheimer disease | EAS | 48 | 2.0 | 1 | 0.938 | 1.000 | 1.067 |
| strict | Alzheimer disease | EUR | 75 | 0.5 | 0 | 0.387 | 0.000 | 0.000 |
| strict | Alzheimer disease | EUR | 75 | 1.0 | 0 | 0.667 | 0.000 | 0.000 |

| | | | | | | | | |
|---|---|---|---|---|---|---|---|---|
| strict | Alzheimer disease | EUR | 75 | 2.0 | 0 | 0.880 | 0.000 | 0.000 |
| strict | Alzheimer disease | SAS | 75 | 0.5 | 0 | 0.333 | 0.000 | 0.000 |
| strict | Alzheimer disease | SAS | 75 | 1.0 | 0 | 0.640 | 0.000 | 0.000 |
| strict | Alzheimer disease | SAS | 75 | 2.0 | 0 | 0.907 | 0.000 | 0.000 |
| strict | LDL cholesterol | AFR | 93 | 0.5 | 0 | 0.344 | 0.000 | 0.000 |
| strict | LDL cholesterol | AFR | 93 | 1.0 | 0 | 0.645 | 0.000 | 0.000 |
| strict | LDL cholesterol | AFR | 93 | 2.0 | 0 | 0.925 | 0.000 | 0.000 |
| strict | LDL cholesterol | EAS | 48 | 0.5 | 0 | 0.354 | 0.000 | 0.000 |
| strict | LDL cholesterol | EAS | 48 | 1.0 | 0 | 0.646 | 0.000 | 0.000 |
| strict | LDL cholesterol | EAS | 48 | 2.0 | 0 | 0.958 | 0.000 | 0.000 |
| strict | LDL cholesterol | EUR | 75 | 0.5 | 0 | 0.387 | 0.000 | 0.000 |
| strict | LDL cholesterol | EUR | 75 | 1.0 | 0 | 0.653 | 0.000 | 0.000 |
| strict | LDL cholester | EUR | 75 | 2.0 | 0 | 0.933 | 0.000 | 0.000 |

| | | | | | | | | |
|---|---|---|---|---|---|---|---|---|
| | ol | | | | | | | |
| strict | LDL cholesterol | SAS | 75 | 0.5 | 0 | 0.373 | 0.000 | 0.000 |
| strict | LDL cholesterol | SAS | 75 | 1.0 | 0 | 0.653 | 0.000 | 0.000 |
| strict | LDL cholesterol | SAS | 75 | 2.0 | 0 | 0.893 | 0.000 | 0.000 |
| strict | Systolic blood pressure | AFR | 93 | 0.5 | 75 | 0.290 | 0.280 | 0.964 |
| strict | Systolic blood pressure | AFR | 93 | 1.0 | 75 | 0.613 | 0.613 | 1.001 |
| strict | Systolic blood pressure | AFR | 93 | 2.0 | 75 | 0.892 | 0.907 | 1.016 |
| strict | Systolic blood pressure | EAS | 48 | 0.5 | 42 | 0.292 | 0.286 | 0.980 |
| strict | Systolic blood pressure | EAS | 48 | 1.0 | 42 | 0.562 | 0.571 | 1.016 |
| strict | Systolic blood pressure | EAS | 48 | 2.0 | 42 | 0.896 | 0.881 | 0.983 |
| strict | Systolic blood pressure | EUR | 75 | 0.5 | 61 | 0.307 | 0.328 | 1.069 |
| strict | Systolic blood pressure | EUR | 75 | 1.0 | 61 | 0.733 | 0.738 | 1.006 |

| | | | | | | | | |
|---|---|---|---|---|---|---|---|---|
| strict | Systolic blood pressure | EUR | 75 | 2.0 | 61 | 0.960 | 0.951 | 0.990 |
| strict | Systolic blood pressure | SAS | 75 | 0.5 | 64 | 0.360 | 0.344 | 0.955 |
| strict | Systolic blood pressure | SAS | 75 | 1.0 | 64 | 0.600 | 0.625 | 1.042 |
| strict | Systolic blood pressure | SAS | 75 | 2.0 | 64 | 0.920 | 0.922 | 1.002 |
| strict | Schizophrenia | AFR | 93 | 0.5 | 0 | 0.366 | 0.000 | 0.000 |
| strict | Schizophrenia | AFR | 93 | 1.0 | 0 | 0.634 | 0.000 | 0.000 |
| strict | Schizophrenia | AFR | 93 | 2.0 | 0 | 0.946 | 0.000 | 0.000 |
| strict | Schizophrenia | EAS | 48 | 0.5 | 0 | 0.250 | 0.000 | 0.000 |
| strict | Schizophrenia | EAS | 48 | 1.0 | 0 | 0.521 | 0.000 | 0.000 |
| strict | Schizophrenia | EAS | 48 | 2.0 | 0 | 0.875 | 0.000 | 0.000 |
| strict | Schizophrenia | EUR | 75 | 0.5 | 0 | 0.307 | 0.000 | 0.000 |
| strict | Schizophrenia | EUR | 75 | 1.0 | 0 | 0.720 | 0.000 | 0.000 |
| strict | Schizophrenia | EUR | 75 | 2.0 | 0 | 0.947 | 0.000 | 0.000 |
| strict | Schizophrenia | SAS | 75 | 0.5 | 0 | 0.347 | 0.000 | 0.000 |
| strict | Schizoph | SAS | 75 | 1.0 | 0 | 0.573 | 0.000 | 0.0 |

| | | | | | | | | |
|---|---|---|---|---|---|---|---|---|
| | renia | | | | | | | 00 |
| strict | Schizophrenia | SAS | 75 | 2.0 | 0 | 0.853 | 0.000 | 0.000 |
| strict | Educational attainment | AFR | 93 | 0.5 | 10 | 0.333 | 0.300 | 0.900 |
| strict | Educational attainment | AFR | 93 | 1.0 | 10 | 0.581 | 0.500 | 0.861 |
| strict | Educational attainment | AFR | 93 | 2.0 | 10 | 0.860 | 0.900 | 1.046 |
| strict | Educational attainment | EAS | 48 | 0.5 | 5 | 0.292 | 0.400 | 1.371 |
| strict | Educational attainment | EAS | 48 | 1.0 | 5 | 0.583 | 0.600 | 1.029 |
| strict | Educational attainment | EAS | 48 | 2.0 | 5 | 0.854 | 0.800 | 0.937 |
| strict | Educational attainment | EUR | 75 | 0.5 | 21 | 0.267 | 0.143 | 0.536 |
| strict | Educational attainment | EUR | 75 | 1.0 | 21 | 0.493 | 0.429 | 0.869 |
| strict | Educational | EUR | 75 | 2.0 | 21 | 0.867 | 0.857 | 0.9 |

| | | | | | | | | |
|---|---|---|---|---|---|---|---|---|
| | attainment | | | | | | | 89 |
| strict | Educational attainment | SAS | 75 | 0.5 | 12 | 0.253 | 0.417 | 1.645 |
| strict | Educational attainment | SAS | 75 | 1.0 | 12 | 0.600 | 0.667 | 1.111 |
| strict | Educational attainment | SAS | 75 | 2.0 | 12 | 0.960 | 1.000 | 1.042 |
| rank | Height | AFR | 93 | 0.5 | 0 | 0.452 | 0.000 | 0.000 |
| rank | Height | AFR | 93 | 1.0 | 0 | 0.742 | 0.000 | 0.000 |
| rank | Height | AFR | 93 | 2.0 | 0 | 0.957 | 0.000 | 0.000 |
| rank | Height | EAS | 48 | 0.5 | 0 | 0.417 | 0.000 | 0.000 |
| rank | Height | EAS | 48 | 1.0 | 0 | 0.708 | 0.000 | 0.000 |
| rank | Height | EAS | 48 | 2.0 | 0 | 0.958 | 0.000 | 0.000 |
| rank | Height | EUR | 75 | 0.5 | 0 | 0.453 | 0.000 | 0.000 |
| rank | Height | EUR | 75 | 1.0 | 0 | 0.747 | 0.000 | 0.000 |
| rank | Height | EUR | 75 | 2.0 | 0 | 0.973 | 0.000 | 0.000 |
| rank | Height | SAS | 75 | 0.5 | 0 | 0.387 | 0.000 | 0.000 |

| | | | | | | | | |
|---|---|---|---|---|---|---|---|---|
| rank | Height | SAS | 75 | 1.0 | 0 | 0.653 | 0.000 | 0.000 |
| rank | Height | SAS | 75 | 2.0 | 0 | 0.960 | 0.000 | 0.000 |
| rank | BMI | AFR | 93 | 0.5 | 2 | 0.376 | 0.500 | 1.329 |
| rank | BMI | AFR | 93 | 1.0 | 2 | 0.591 | 0.500 | 0.845 |
| rank | BMI | AFR | 93 | 2.0 | 2 | 0.925 | 1.000 | 1.081 |
| rank | BMI | EAS | 48 | 0.5 | 0 | 0.479 | 0.000 | 0.000 |
| rank | BMI | EAS | 48 | 1.0 | 0 | 0.667 | 0.000 | 0.000 |
| rank | BMI | EAS | 48 | 2.0 | 0 | 0.875 | 0.000 | 0.000 |
| rank | BMI | EUR | 75 | 0.5 | 12 | 0.333 | 0.417 | 1.250 |
| rank | BMI | EUR | 75 | 1.0 | 12 | 0.600 | 0.667 | 1.111 |
| rank | BMI | EUR | 75 | 2.0 | 12 | 0.933 | 1.000 | 1.071 |
| rank | BMI | SAS | 75 | 0.5 | 9 | 0.440 | 0.444 | 1.010 |
| rank | BMI | SAS | 75 | 1.0 | 9 | 0.733 | 0.444 | 0.606 |
| rank | BMI | SAS | 75 | 2.0 | 9 | 0.947 | 0.778 | 0.822 |
| rank | T2D | AFR | 93 | 0.5 | 0 | 0.258 | 0.000 | 0.000 |
| rank | T2D | AFR | 93 | 1.0 | 0 | 0.559 | 0.000 | 0.000 |

| | | | | | | | | |
|---|---|---|---|---|---|---|---|---|
| rank | T2D | AFR | 93 | 2.0 | 0 | 0.892 | 0.000 | 0.000 |
| rank | T2D | EAS | 48 | 0.5 | 1 | 0.417 | 0.000 | 0.000 |
| rank | T2D | EAS | 48 | 1.0 | 1 | 0.667 | 0.000 | 0.000 |
| rank | T2D | EAS | 48 | 2.0 | 1 | 0.833 | 0.000 | 0.000 |
| rank | T2D | EUR | 75 | 0.5 | 15 | 0.227 | 0.200 | 0.882 |
| rank | T2D | EUR | 75 | 1.0 | 15 | 0.493 | 0.400 | 0.811 |
| rank | T2D | EUR | 75 | 2.0 | 15 | 0.800 | 0.800 | 1.000 |
| rank | T2D | SAS | 75 | 0.5 | 3 | 0.280 | 0.000 | 0.000 |
| rank | T2D | SAS | 75 | 1.0 | 3 | 0.573 | 0.333 | 0.581 |
| rank | T2D | SAS | 75 | 2.0 | 3 | 0.880 | 0.667 | 0.758 |
| rank | CAD | AFR | 93 | 0.5 | 0 | 0.290 | 0.000 | 0.000 |
| rank | CAD | AFR | 93 | 1.0 | 0 | 0.591 | 0.000 | 0.000 |
| rank | CAD | AFR | 93 | 2.0 | 0 | 0.925 | 0.000 | 0.000 |
| rank | CAD | EAS | 48 | 0.5 | 0 | 0.229 | 0.000 | 0.000 |
| rank | CAD | EAS | 48 | 1.0 | 0 | 0.625 | 0.000 | 0.000 |
| rank | CAD | EAS | 48 | 2.0 | 0 | 0.875 | 0.000 | 0.000 |

| | | | | | | | | |
|---|---|---|---|---|---|---|---|---|
| rank | CAD | EUR | 75 | 0.5 | 0 | 0.307 | 0.000 | 0.0<br>00 |
| rank | CAD | EUR | 75 | 1.0 | 0 | 0.533 | 0.000 | 0.0<br>00 |
| rank | CAD | EUR | 75 | 2.0 | 0 | 0.867 | 0.000 | 0.0<br>00 |
| rank | CAD | SAS | 75 | 0.5 | 0 | 0.320 | 0.000 | 0.0<br>00 |
| rank | CAD | SAS | 75 | 1.0 | 0 | 0.547 | 0.000 | 0.0<br>00 |
| rank | CAD | SAS | 75 | 2.0 | 0 | 0.867 | 0.000 | 0.0<br>00 |
| rank | Breast cancer | AFR | 93 | 0.5 | 17 | 0.409 | 0.294 | 0.7<br>20 |
| rank | Breast cancer | AFR | 93 | 1.0 | 17 | 0.624 | 0.588 | 0.9<br>43 |
| rank | Breast cancer | AFR | 93 | 2.0 | 17 | 0.871 | 0.882 | 1.0<br>13 |
| rank | Breast cancer | EAS | 48 | 0.5 | 5 | 0.292 | 0.400 | 1.3<br>71 |
| rank | Breast cancer | EAS | 48 | 1.0 | 5 | 0.583 | 0.800 | 1.3<br>71 |
| rank | Breast cancer | EAS | 48 | 2.0 | 5 | 0.917 | 1.000 | 1.0<br>91 |
| rank | Breast cancer | EUR | 75 | 0.5 | 11 | 0.320 | 0.545 | 1.7<br>05 |
| rank | Breast cancer | EUR | 75 | 1.0 | 11 | 0.627 | 0.636 | 1.0<br>15 |
| rank | Breast cancer | EUR | 75 | 2.0 | 11 | 0.933 | 0.909 | 0.9<br>74 |
| rank | Breast cancer | SAS | 75 | 0.5 | 10 | 0.280 | 0.400 | 1.4<br>29 |

| | | | | | | | | |
|---|---|---|---|---|---|---|---|---|
| rank | Breast cancer | SAS | 75 | 1.0 | 10 | 0.547 | 0.600 | 1.098 |
| rank | Breast cancer | SAS | 75 | 2.0 | 10 | 0.880 | 1.000 | 1.136 |
| rank | Alzheimer disease | AFR | 93 | 0.5 | 14 | 0.301 | 0.143 | 0.474 |
| rank | Alzheimer disease | AFR | 93 | 1.0 | 14 | 0.591 | 0.286 | 0.483 |
| rank | Alzheimer disease | AFR | 93 | 2.0 | 14 | 0.903 | 0.857 | 0.949 |
| rank | Alzheimer disease | EAS | 48 | 0.5 | 34 | 0.229 | 0.235 | 1.027 |
| rank | Alzheimer disease | EAS | 48 | 1.0 | 34 | 0.479 | 0.500 | 1.043 |
| rank | Alzheimer disease | EAS | 48 | 2.0 | 34 | 0.938 | 0.971 | 1.035 |
| rank | Alzheimer disease | EUR | 75 | 0.5 | 56 | 0.387 | 0.446 | 1.155 |
| rank | Alzheimer disease | EUR | 75 | 1.0 | 56 | 0.667 | 0.732 | 1.098 |
| rank | Alzheimer disease | EUR | 75 | 2.0 | 56 | 0.880 | 0.911 | 1.035 |
| rank | Alzheimer disease | SAS | 75 | 0.5 | 51 | 0.333 | 0.412 | 1.235 |
| rank | Alzheimer disease | SAS | 75 | 1.0 | 51 | 0.640 | 0.706 | 1.103 |
| rank | Alzheimer disease | SAS | 75 | 2.0 | 51 | 0.907 | 0.941 | 1.038 |
| rank | LDL cholesterol | AFR | 93 | 0.5 | 5 | 0.344 | 0.400 | 1.162 |
| rank | LDL cholester | AFR | 93 | 1.0 | 5 | 0.645 | 0.600 | 0.9 |

| | | | | | | | | |
|---|---|---|---|---|---|---|---|---|
| | ol | | | | | | | 30 |
| rank | LDL cholesterol | AFR | 93 | 2.0 | 5 | 0.925 | 0.800 | 0.865 |
| rank | LDL cholesterol | EAS | 48 | 0.5 | 23 | 0.354 | 0.304 | 0.859 |
| rank | LDL cholesterol | EAS | 48 | 1.0 | 23 | 0.646 | 0.565 | 0.875 |
| rank | LDL cholesterol | EAS | 48 | 2.0 | 23 | 0.958 | 1.000 | 1.043 |
| rank | LDL cholesterol | EUR | 75 | 0.5 | 22 | 0.387 | 0.455 | 1.176 |
| rank | LDL cholesterol | EUR | 75 | 1.0 | 22 | 0.653 | 0.727 | 1.113 |
| rank | LDL cholesterol | EUR | 75 | 2.0 | 22 | 0.933 | 0.955 | 1.023 |
| rank | LDL cholesterol | SAS | 75 | 0.5 | 28 | 0.373 | 0.357 | 0.957 |
| rank | LDL cholesterol | SAS | 75 | 1.0 | 28 | 0.653 | 0.607 | 0.929 |
| rank | LDL cholesterol | SAS | 75 | 2.0 | 28 | 0.893 | 0.929 | 1.039 |
| rank | Systolic blood pressure | AFR | 93 | 0.5 | 19 | 0.290 | 0.158 | 0.544 |

| | | | | | | | | |
|---|---|---|---|---|---|---|---|---|
| rank | Systolic blood pressure | AFR | 93 | 1.0 | 19 | 0.613 | 0.368 | 0.601 |
| rank | Systolic blood pressure | AFR | 93 | 2.0 | 19 | 0.892 | 0.895 | 1.003 |
| rank | Systolic blood pressure | EAS | 48 | 0.5 | 30 | 0.292 | 0.267 | 0.914 |
| rank | Systolic blood pressure | EAS | 48 | 1.0 | 30 | 0.562 | 0.433 | 0.770 |
| rank | Systolic blood pressure | EAS | 48 | 2.0 | 30 | 0.896 | 0.867 | 0.967 |
| rank | Systolic blood pressure | EUR | 75 | 0.5 | 38 | 0.307 | 0.395 | 1.287 |
| rank | Systolic blood pressure | EUR | 75 | 1.0 | 38 | 0.733 | 0.763 | 1.041 |
| rank | Systolic blood pressure | EUR | 75 | 2.0 | 38 | 0.960 | 0.947 | 0.987 |
| rank | Systolic blood pressure | SAS | 75 | 0.5 | 38 | 0.360 | 0.421 | 1.170 |
| rank | Systolic blood pressure | SAS | 75 | 1.0 | 38 | 0.600 | 0.658 | 1.096 |
| rank | Systolic blood pressure | SAS | 75 | 2.0 | 38 | 0.920 | 0.921 | 1.001 |
| rank | Schizophrenia | AFR | 93 | 0.5 | 0 | 0.366 | 0.000 | 0.000 |

| | | | | | | | | |
|---|---|---|---|---|---|---|---|---|
| rank | Schizophrenia | AFR | 93 | 1.0 | 0 | 0.634 | 0.000 | 0.000 |
| rank | Schizophrenia | AFR | 93 | 2.0 | 0 | 0.946 | 0.000 | 0.000 |
| rank | Schizophrenia | EAS | 48 | 0.5 | 0 | 0.250 | 0.000 | 0.000 |
| rank | Schizophrenia | EAS | 48 | 1.0 | 0 | 0.521 | 0.000 | 0.000 |
| rank | Schizophrenia | EAS | 48 | 2.0 | 0 | 0.875 | 0.000 | 0.000 |
| rank | Schizophrenia | EUR | 75 | 0.5 | 0 | 0.307 | 0.000 | 0.000 |
| rank | Schizophrenia | EUR | 75 | 1.0 | 0 | 0.720 | 0.000 | 0.000 |
| rank | Schizophrenia | EUR | 75 | 2.0 | 0 | 0.947 | 0.000 | 0.000 |
| rank | Schizophrenia | SAS | 75 | 0.5 | 0 | 0.347 | 0.000 | 0.000 |
| rank | Schizophrenia | SAS | 75 | 1.0 | 0 | 0.573 | 0.000 | 0.000 |
| rank | Schizophrenia | SAS | 75 | 2.0 | 0 | 0.853 | 0.000 | 0.000 |
| rank | Educational attainment | AFR | 93 | 0.5 | 0 | 0.333 | 0.000 | 0.000 |
| rank | Educational attainment | AFR | 93 | 1.0 | 0 | 0.581 | 0.000 | 0.000 |
| rank | Educational attainment | AFR | 93 | 2.0 | 0 | 0.860 | 0.000 | 0.000 |

| | | | | | | | | |
|---|---|---|---|---|---|---|---|---|
| rank | Educational attainment | EAS | 48 | 0.5 | 0 | 0.292 | 0.000 | 0.000 |
| rank | Educational attainment | EAS | 48 | 1.0 | 0 | 0.583 | 0.000 | 0.000 |
| rank | Educational attainment | EAS | 48 | 2.0 | 0 | 0.854 | 0.000 | 0.000 |
| rank | Educational attainment | EUR | 75 | 0.5 | 0 | 0.267 | 0.000 | 0.000 |
| rank | Educational attainment | EUR | 75 | 1.0 | 0 | 0.493 | 0.000 | 0.000 |
| rank | Educational attainment | EUR | 75 | 2.0 | 0 | 0.867 | 0.000 | 0.000 |
| rank | Educational attainment | SAS | 75 | 0.5 | 0 | 0.253 | 0.000 | 0.000 |
| rank | Educational attainment | SAS | 75 | 1.0 | 0 | 0.600 | 0.000 | 0.000 |
| rank | Educational attainment | SAS | 75 | 2.0 | 0 | 0.960 | 0.000 | 0.000 |
| reca | Height | AFR | 93 | 0.5 | 37 | 0.452 | 0.432 | 0.9 |

| | | | | | | | | |
|---|---|---|---|---|---|---|---|---|
| l | | | | | | | | 58 |
| recal | Height | AFR | 93 | 1.0 | 37 | 0.742 | 0.730 | 0.984 |
| recal | Height | AFR | 93 | 2.0 | 37 | 0.957 | 1.000 | 1.045 |
| recal | Height | EAS | 48 | 0.5 | 4 | 0.417 | 0.250 | 0.600 |
| recal | Height | EAS | 48 | 1.0 | 4 | 0.708 | 0.250 | 0.353 |
| recal | Height | EAS | 48 | 2.0 | 4 | 0.958 | 0.750 | 0.783 |
| recal | Height | EUR | 75 | 0.5 | 7 | 0.453 | 0.571 | 1.261 |
| recal | Height | EUR | 75 | 1.0 | 7 | 0.747 | 0.857 | 1.148 |
| recal | Height | EUR | 75 | 2.0 | 7 | 0.973 | 1.000 | 1.027 |
| recal | Height | SAS | 75 | 0.5 | 0 | 0.387 | 0.000 | 0.000 |
| recal | Height | SAS | 75 | 1.0 | 0 | 0.653 | 0.000 | 0.000 |
| recal | Height | SAS | 75 | 2.0 | 0 | 0.960 | 0.000 | 0.000 |
| recal | BMI | AFR | 93 | 0.5 | 2 | 0.376 | 0.000 | 0.000 |
| recal | BMI | AFR | 93 | 1.0 | 2 | 0.591 | 0.500 | 0.845 |
| recal | BMI | AFR | 93 | 2.0 | 2 | 0.925 | 1.000 | 1.081 |
| recal | BMI | EAS | 48 | 0.5 | 2 | 0.479 | 0.500 | 1.043 |

| | | | | | | | | |
|---|---|---|---|---|---|---|---|---|
| recal | BMI | EAS | 48 | 1.0 | 2 | 0.667 | 0.500 | 0.750 |
| recal | BMI | EAS | 48 | 2.0 | 2 | 0.875 | 1.000 | 1.143 |
| recal | BMI | EUR | 75 | 0.5 | 7 | 0.333 | 0.429 | 1.286 |
| recal | BMI | EUR | 75 | 1.0 | 7 | 0.600 | 0.714 | 1.190 |
| recal | BMI | EUR | 75 | 2.0 | 7 | 0.933 | 0.857 | 0.918 |
| recal | BMI | SAS | 75 | 0.5 | 5 | 0.440 | 0.400 | 0.909 |
| recal | BMI | SAS | 75 | 1.0 | 5 | 0.733 | 0.600 | 0.818 |
| recal | BMI | SAS | 75 | 2.0 | 5 | 0.947 | 1.000 | 1.056 |
| recal | T2D | AFR | 93 | 0.5 | 0 | 0.258 | 0.000 | 0.000 |
| recal | T2D | AFR | 93 | 1.0 | 0 | 0.559 | 0.000 | 0.000 |
| recal | T2D | AFR | 93 | 2.0 | 0 | 0.892 | 0.000 | 0.000 |
| recal | T2D | EAS | 48 | 0.5 | 0 | 0.417 | 0.000 | 0.000 |
| recal | T2D | EAS | 48 | 1.0 | 0 | 0.667 | 0.000 | 0.000 |
| recal | T2D | EAS | 48 | 2.0 | 0 | 0.833 | 0.000 | 0.000 |
| recal | T2D | EUR | 75 | 0.5 | 6 | 0.227 | 0.167 | 0.735 |
| recal | T2D | EUR | 75 | 1.0 | 6 | 0.493 | 0.167 | 0.338 |

| | | | | | | | | |
|---|---|---|---|---|---|---|---|---|
| recal | T2D | EUR | 75 | 2.0 | 6 | 0.800 | 0.667 | 0.833 |
| recal | T2D | SAS | 75 | 0.5 | 2 | 0.280 | 0.500 | 1.786 |
| recal | T2D | SAS | 75 | 1.0 | 2 | 0.573 | 1.000 | 1.744 |
| recal | T2D | SAS | 75 | 2.0 | 2 | 0.880 | 1.000 | 1.136 |
| recal | CAD | AFR | 93 | 0.5 | 12 | 0.290 | 0.500 | 1.722 |
| recal | CAD | AFR | 93 | 1.0 | 12 | 0.591 | 0.750 | 1.268 |
| recal | CAD | AFR | 93 | 2.0 | 12 | 0.925 | 1.000 | 1.081 |
| recal | CAD | EAS | 48 | 0.5 | 1 | 0.229 | 0.000 | 0.000 |
| recal | CAD | EAS | 48 | 1.0 | 1 | 0.625 | 0.000 | 0.000 |
| recal | CAD | EAS | 48 | 2.0 | 1 | 0.875 | 0.000 | 0.000 |
| recal | CAD | EUR | 75 | 0.5 | 11 | 0.307 | 0.091 | 0.296 |
| recal | CAD | EUR | 75 | 1.0 | 11 | 0.533 | 0.636 | 1.193 |
| recal | CAD | EUR | 75 | 2.0 | 11 | 0.867 | 1.000 | 1.154 |
| recal | CAD | SAS | 75 | 0.5 | 5 | 0.320 | 0.200 | 0.625 |
| recal | CAD | SAS | 75 | 1.0 | 5 | 0.547 | 0.200 | 0.366 |
| recal | CAD | SAS | 75 | 2.0 | 5 | 0.867 | 0.600 | 0.692 |

| | | | | | | | | |
|---|---|---|---|---|---|---|---|---|
| recal | Breast cancer | AFR | 93 | 0.5 | 26 | 0.409 | 0.385 | 0.941 |
| recal | Breast cancer | AFR | 93 | 1.0 | 26 | 0.624 | 0.577 | 0.925 |
| recal | Breast cancer | AFR | 93 | 2.0 | 26 | 0.871 | 0.808 | 0.927 |
| recal | Breast cancer | EAS | 48 | 0.5 | 5 | 0.292 | 0.400 | 1.371 |
| recal | Breast cancer | EAS | 48 | 1.0 | 5 | 0.583 | 0.800 | 1.371 |
| recal | Breast cancer | EAS | 48 | 2.0 | 5 | 0.917 | 1.000 | 1.091 |
| recal | Breast cancer | EUR | 75 | 0.5 | 7 | 0.320 | 0.429 | 1.339 |
| recal | Breast cancer | EUR | 75 | 1.0 | 7 | 0.627 | 0.571 | 0.912 |
| recal | Breast cancer | EUR | 75 | 2.0 | 7 | 0.933 | 0.857 | 0.918 |
| recal | Breast cancer | SAS | 75 | 0.5 | 10 | 0.280 | 0.200 | 0.714 |
| recal | Breast cancer | SAS | 75 | 1.0 | 10 | 0.547 | 0.400 | 0.732 |
| recal | Breast cancer | SAS | 75 | 2.0 | 10 | 0.880 | 0.800 | 0.909 |
| recal | Alzheimer disease | AFR | 93 | 0.5 | 5 | 0.301 | 0.200 | 0.664 |
| recal | Alzheimer disease | AFR | 93 | 1.0 | 5 | 0.591 | 0.400 | 0.676 |
| recal | Alzheimer disease | AFR | 93 | 2.0 | 5 | 0.903 | 0.800 | 0.886 |
| recal | Alzheimer disease | EAS | 48 | 0.5 | 4 | 0.229 | 0.500 | 2.182 |

| | | | | | | | | |
|---|---|---|---|---|---|---|---|---|
| recall | Alzheimer disease | EAS | 48 | 1.0 | 4 | 0.479 | 1.000 | 2.087 |
| recall | Alzheimer disease | EAS | 48 | 2.0 | 4 | 0.938 | 1.000 | 1.067 |
| recall | Alzheimer disease | EUR | 75 | 0.5 | 5 | 0.387 | 0.600 | 1.552 |
| recall | Alzheimer disease | EUR | 75 | 1.0 | 5 | 0.667 | 1.000 | 1.500 |
| recall | Alzheimer disease | EUR | 75 | 2.0 | 5 | 0.880 | 1.000 | 1.136 |
| recall | Alzheimer disease | SAS | 75 | 0.5 | 10 | 0.333 | 0.200 | 0.600 |
| recall | Alzheimer disease | SAS | 75 | 1.0 | 10 | 0.640 | 0.500 | 0.781 |
| recall | Alzheimer disease | SAS | 75 | 2.0 | 10 | 0.907 | 1.000 | 1.103 |
| recall | LDL cholesterol | AFR | 93 | 0.5 | 0 | 0.344 | 0.000 | 0.000 |
| recall | LDL cholesterol | AFR | 93 | 1.0 | 0 | 0.645 | 0.000 | 0.000 |
| recall | LDL cholesterol | AFR | 93 | 2.0 | 0 | 0.925 | 0.000 | 0.000 |
| recall | LDL cholesterol | EAS | 48 | 0.5 | 8 | 0.354 | 0.375 | 1.059 |
| recall | LDL cholesterol | EAS | 48 | 1.0 | 8 | 0.646 | 0.875 | 1.355 |
| recall | LDL cholesterol | EAS | 48 | 2.0 | 8 | 0.958 | 1.000 | 1.043 |

| | | | | | | | | |
|---|---|---|---|---|---|---|---|---|
| recal | LDL cholesterol | EUR | 75 | 0.5 | 9 | 0.387 | 0.444 | 1.149 |
| recal | LDL cholesterol | EUR | 75 | 1.0 | 9 | 0.653 | 0.889 | 1.361 |
| recal | LDL cholesterol | EUR | 75 | 2.0 | 9 | 0.933 | 1.000 | 1.071 |
| recal | LDL cholesterol | SAS | 75 | 0.5 | 6 | 0.373 | 0.500 | 1.339 |
| recal | LDL cholesterol | SAS | 75 | 1.0 | 6 | 0.653 | 0.667 | 1.020 |
| recal | LDL cholesterol | SAS | 75 | 2.0 | 6 | 0.893 | 1.000 | 1.119 |
| recal | Systolic blood pressure | AFR | 93 | 0.5 | 2 | 0.290 | 0.000 | 0.000 |
| recal | Systolic blood pressure | AFR | 93 | 1.0 | 2 | 0.613 | 0.500 | 0.816 |
| recal | Systolic blood pressure | AFR | 93 | 2.0 | 2 | 0.892 | 1.000 | 1.120 |
| recal | Systolic blood pressure | EAS | 48 | 0.5 | 5 | 0.292 | 0.000 | 0.000 |
| recal | Systolic blood pressure | EAS | 48 | 1.0 | 5 | 0.562 | 0.200 | 0.356 |
| recal | Systolic blood | EAS | 48 | 2.0 | 5 | 0.896 | 0.800 | 0.893 |

| | | | | | | | | |
|---|---|---|---|---|---|---|---|---|
| | pressure | | | | | | | |
| recal | Systolic blood pressure | EUR | 75 | 0.5 | 4 | 0.307 | 0.500 | 1.630 |
| recal | Systolic blood pressure | EUR | 75 | 1.0 | 4 | 0.733 | 0.500 | 0.682 |
| recal | Systolic blood pressure | EUR | 75 | 2.0 | 4 | 0.960 | 1.000 | 1.042 |
| recal | Systolic blood pressure | SAS | 75 | 0.5 | 3 | 0.360 | 1.000 | 2.778 |
| recal | Systolic blood pressure | SAS | 75 | 1.0 | 3 | 0.600 | 1.000 | 1.667 |
| recal | Systolic blood pressure | SAS | 75 | 2.0 | 3 | 0.920 | 1.000 | 1.087 |
| recal | Schizophrenia | AFR | 93 | 0.5 | 0 | 0.366 | 0.000 | 0.000 |
| recal | Schizophrenia | AFR | 93 | 1.0 | 0 | 0.634 | 0.000 | 0.000 |
| recal | Schizophrenia | AFR | 93 | 2.0 | 0 | 0.946 | 0.000 | 0.000 |
| recal | Schizophrenia | EAS | 48 | 0.5 | 0 | 0.250 | 0.000 | 0.000 |
| recal | Schizophrenia | EAS | 48 | 1.0 | 0 | 0.521 | 0.000 | 0.000 |
| recal | Schizophrenia | EAS | 48 | 2.0 | 0 | 0.875 | 0.000 | 0.000 |
| recal | Schizophrenia | EUR | 75 | 0.5 | 10 | 0.307 | 0.300 | 0.978 |

| | | | | | | | | |
|---|---|---|---|---|---|---|---|---|
| recall | Schizophrenia | EUR | 75 | 1.0 | 10 | 0.720 | 0.800 | 1.111 |
| recall | Schizophrenia | EUR | 75 | 2.0 | 10 | 0.947 | 1.000 | 1.056 |
| recall | Schizophrenia | SAS | 75 | 0.5 | 1 | 0.347 | 1.000 | 2.885 |
| recall | Schizophrenia | SAS | 75 | 1.0 | 1 | 0.573 | 1.000 | 1.744 |
| recall | Schizophrenia | SAS | 75 | 2.0 | 1 | 0.853 | 1.000 | 1.172 |
| recall | Educational attainment | AFR | 93 | 0.5 | 2 | 0.333 | 0.000 | 0.000 |
| recall | Educational attainment | AFR | 93 | 1.0 | 2 | 0.581 | 0.500 | 0.861 |
| recall | Educational attainment | AFR | 93 | 2.0 | 2 | 0.860 | 0.500 | 0.581 |
| recall | Educational attainment | EAS | 48 | 0.5 | 0 | 0.292 | 0.000 | 0.000 |
| recall | Educational attainment | EAS | 48 | 1.0 | 0 | 0.583 | 0.000 | 0.000 |
| recall | Educational attainment | EAS | 48 | 2.0 | 0 | 0.854 | 0.000 | 0.000 |
| reca | Educational | EUR | 75 | 0.5 | 11 | 0.267 | 0.091 | 0.3 |

| | | | | | | | | |
|---|---|---|---|---|---|---|---|---|
| l | attainment | | | | | | | 41 |
| recall | Educational attainment | EUR | 75 | 1.0 | 11 | 0.493 | 0.182 | 0.369 |
| recall | Educational attainment | EUR | 75 | 2.0 | 11 | 0.867 | 0.727 | 0.839 |
| recall | Educational attainment | SAS | 75 | 0.5 | 3 | 0.253 | 0.667 | 2.632 |
| recall | Educational attainment | SAS | 75 | 1.0 | 3 | 0.600 | 0.667 | 1.111 |
| recall | Educational attainment | SAS | 75 | 2.0 | 3 | 0.960 | 1.000 | 1.042 |

### S3. Supplementary figures

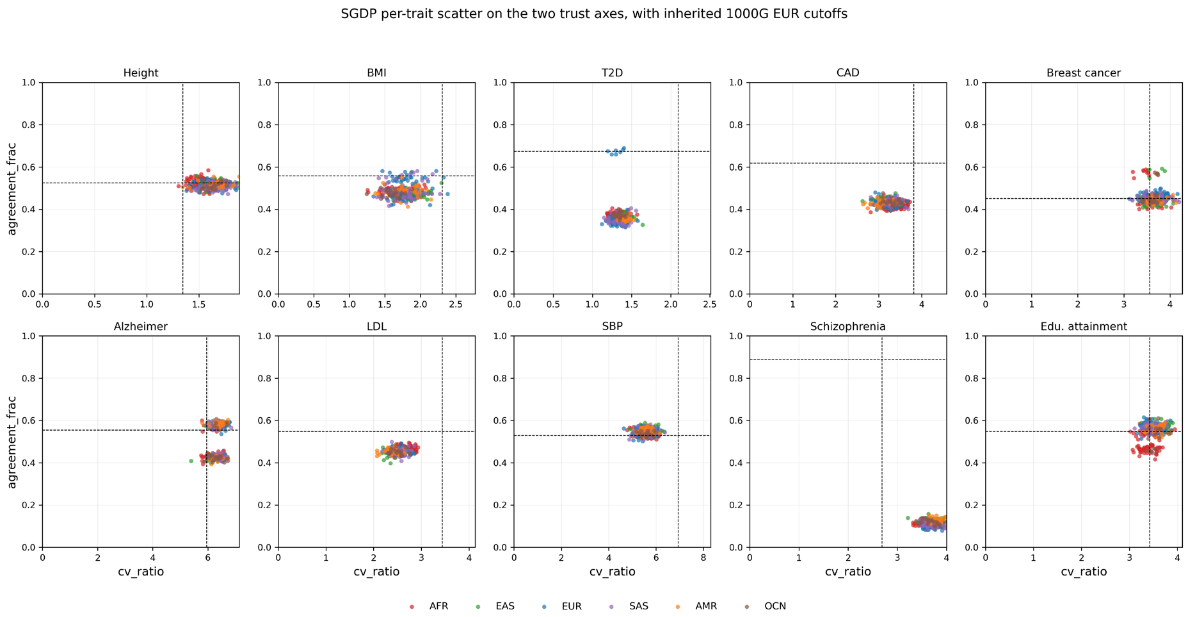


**Figure S1.** Per-trait scatter of cv_ratio (x) and agreement_frac (y) on SGDP individuals coloured by approximate 1000G super-population, with the inherited 1000G EUR cutoffs (regime 1, strict OOD) shown as horizontal and vertical dashed lines. The four quadrants formed by the cutoffs map to scenarios A (top-left), B (bottom-left), C (top-right), and D (bottom-right). Schizophrenia (bottom row, fourth panel) is the cleanest example of distribution-shape mismatch, with the entire SGDP cluster sitting at agreement_frac approximately 0.1 while the EUR cutoff sits at approximately 0.9; no monotone rescaling of the agreement axis can place those points on the trustworthy side of the cutoff. Super-population assignments for SGDP are derived from a hardcoded population-to-region map based on Mallick et al. 2016 because the canonical SGDP metadata table is on the HPC scratch filesystem and is not bundled with the released code; a few populations near regional boundaries (e.g. Tu, Mongola, Yi) may shift one cell in the legend without affecting the per-trait collapse pattern that the figure communicates.

### S4. Supplementary tables

Table S2 provides full enrichment factor values at thresholds of 0.5, 1.0, and 2.0 SD for all three calibration regimes, all ten traits, and four SGDP super-populations. It is available in the supplementary data archive as a tab-separated file and as a formatted table with caption.

Table S1 provides the provenance of the two cross-cohort corrections described in S2, including the symptom, the affected output, and the remedy for each. It is available in the supplementary data archive as a tab-separated file and as a formatted table with caption.

### S5. Data availability

Source data. 1000G phase-3 release files are downloaded from ftp.1000genomes.ebi.ac.uk; SBayesRC 115-trait weight files are downloaded from gctbhub.cloud.edu.au/data/SBayesRC/share/115traits; SGDP cteam_extended.v4.maf0.1perc files are downloaded from sharehost.hms.harvard.edu under the SGDP public release. All sources are free for academic use; the SGDP signed-letter samples (n=21) are not used in this work.
